# Increased burden of rare risk variants across gene expression networks predisposes to sporadic Parkinson’s disease

**DOI:** 10.1101/2024.08.30.610195

**Authors:** Elena Eubanks, Katelyn VanderSleen, Jiya Mody, Neha Patel, Benjamin Sacks, Mahsa Darestani Farahani, Jinying Wang, Jordan Elliott, Nora Jaber, Fulya Akçimen, Sara Bandres-Ciga, Fadel Helweh, Jun Liu, Sanjana Archakam, Robert Kimelman, Bineet Sharma, Philip Socha, Ananya Guntur, Tim Bartels, Ulf Dettmer, M. Maral Mouradian, Amir Houshang Bahrami, Wei Dai, Jean Baum, Zheng Shi, John Hardy, Eleanna Kara

## Abstract

Alpha-synuclein (αSyn) is an intrinsically disordered protein that accumulates in the brains of patients with Parkinson’s disease and forms intraneuronal inclusions called Lewy Bodies. While the mechanism underlying the dysregulation of αSyn in Parkinson’s disease is unclear, it is thought that prionoid cell-to-cell propagation of αSyn has an important role. Through a high throughput screen, we recently identified 38 genes whose knock down modulates αSyn propagation. Follow up experiments were undertaken for two of those genes, *TAX1BP1* and *ADAMTS19*, to study the mechanism with which they regulate αSyn homeostasis. We used a recently developed M17D neuroblastoma cell line expressing triple mutant (E35K+E46K+E61K) “3K” αSyn under doxycycline induction. 3K αSyn spontaneously forms inclusions that show ultrastructural similarities to Lewy Bodies. Experiments using that cell line showed that *TAX1BP1* and *ADAMTS19* regulate how αSyn interacts with lipids and phase separates into inclusions, respectively, adding to the growing body of evidence implicating those processes in Parkinson’s disease. Through RNA sequencing, we identified several genes that are differentially expressed after knock-down of *TAX1BP1* or *ADAMTS19*. Burden analysis revealed that those differentially expressed genes (DEGs) carry an increased frequency of rare risk variants in Parkinson’s disease patients versus healthy controls, an effect that was independently replicated across two separate cohorts (GP2 and AMP-PD). Weighted gene co-expression network analysis (WGCNA) showed that the DEGs cluster within modules in regions of the brain that develop high degrees of αSyn pathology (basal ganglia, cortex). We propose a novel model for the genetic architecture of sporadic Parkinson’s disease: increased burden of risk variants across genetic networks dysregulates pathways underlying αSyn homeostasis, thereby leading to pathology and neurodegeneration.

## INTRODUCTION

Alpha-synuclein (αSyn) is an intrinsically disordered protein with a critical role in the pathogenesis of Parkinson’s disease (PD), Dementia with Lewy Bodies (DLB) and multiple system atrophy (MSA). Point mutations and whole gene multiplications in the *SNCA* gene cause rare Mendelian forms of PD. Its N-terminus and NAC domain (fig.1A) contain nine imperfect KTKEGV amino acid repeat motifs that mediate its membrane binding properties. Specifically, the repeat motifs underlie the formation of amphipathic helices that facilitate membrane binding; the positively charged lysines (K) within the repeat motifs interact with negatively charged lipid headgroups (Jao *et al*, 2008), while the non-polar amino acids in the hydrophobic region of the protein insert below the headgroups and interact with lipid tails through van der Waals forces (Tsigelny *et al*, 2012; Wietek *et al*, 2013).

**Figure 1:**
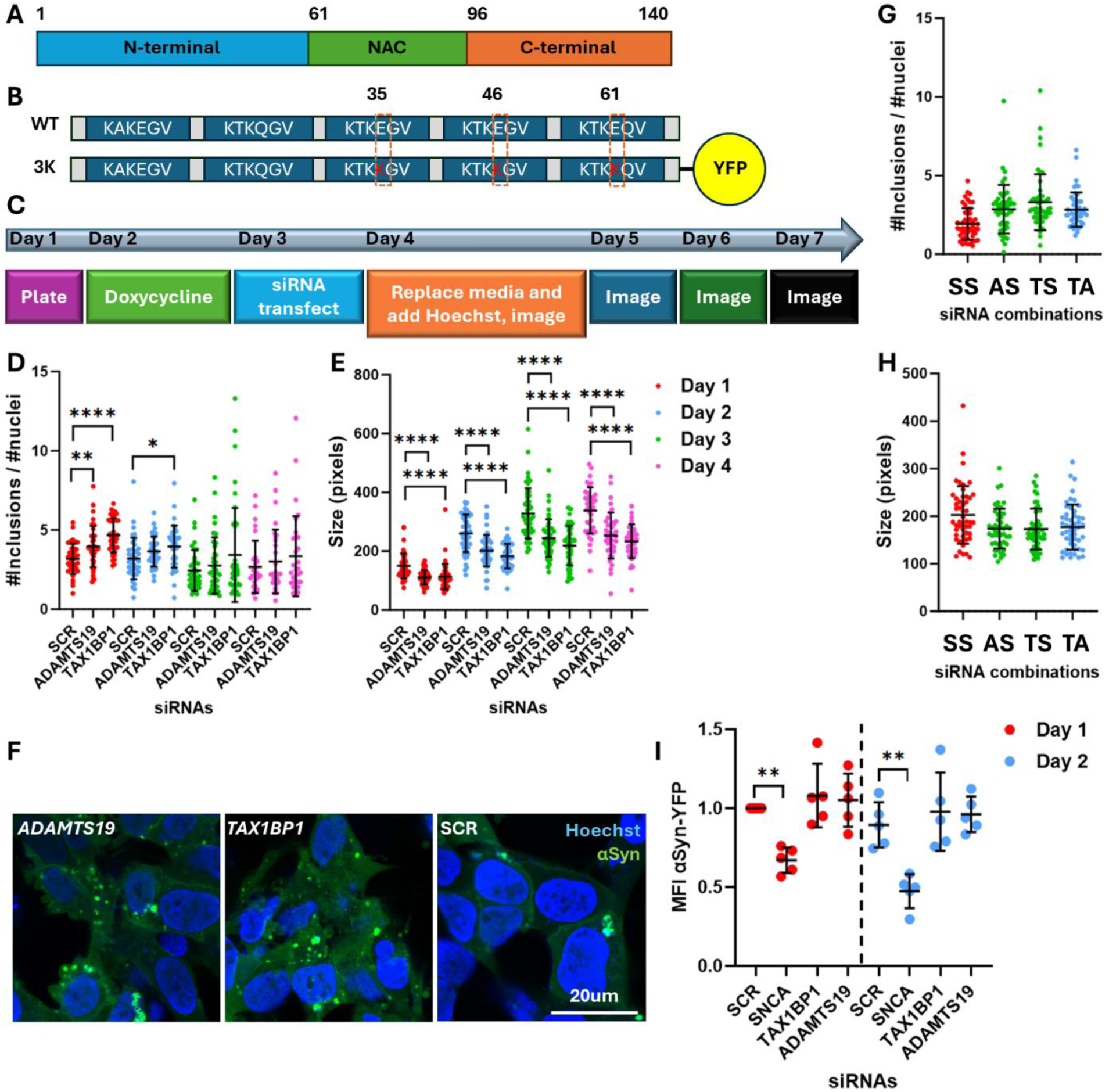
Effect of *TAX1BP1* or *ADAMTS19* knock down on αSyn inclusions. A. Diagram showing the regions within the αSyn protein: N-terminal domain, non-Aβ component (NAC), C-terminal acidic tail. B. Diagram showing the location of the mutations in the 3K αSyn mutant protein. The E46K is a naturally occurring Mendelian mutation, whereas the E35K and E61K are artificial mutations in the flanking repeat motifs. C. Workflow of the experiments undertaken on the 3K αSyn cell line: Cells were plated on day 1, treated with doxycycline on day 2, transfected with siRNAs on day 3, fed with medium containing no phenol red on day 4, and then live imaged on day 4 onwards. D. Effect of knock-down of *TAX1BP1* and *ADAMTS19* versus scrambled (SCR) on the number and E. size of αSyn inclusions over several days. One way ANOVA with Dunnett test correction for multiple testing. F. Representative confocal images showing the effect of the knock-downs on αSyn inclusions. G. Effect of the combined effect of *TAX1BP1* and *ADAMTS19* knock-down on the number and H. size of αSyn inclusions. X axis labelling refers to the co-transfection combinations: AS: *ADAMTS19*+SCR+SCR; TS: *TAX1BP1*+SCR+SCR; TA: *TAX1BP1*, *ADAMTS19*, SCR. One way ANOVA with Dunnett test correction for multiple testing. I. Flow cytometry plots at days 1 and 2 post-transfection showing the effect of the *TAX1BP1* and *ADAMTS19* knock-downs on the amount of αSyn per cell, as inferred by the mean fluorescence intensity (MFI) of the YFP tag. The positive control, siRNA targeting *SNCA*, showed a significant reduction in MFI for YFP on both days. One way ANOVA with Dunnett test correction for multiple testing (undertaken separately for each day). 5 biological (independent) replicates were done for each experiment, unless otherwise stated. P values: *≤0.05, **≤0.01, ***≤0.001, ****≤0.0001. Only statistically significant differences are shown in the plots. Abbreviations: SCR: scrambled.

Altered interactions between αSyn and membranes are thought to underlie the process with which αSyn dysregulation in PD leads to the formation of Lewy Bodies (Kim *et al*, 2021a). Recent quantitative correlative light and electron microscopy (CLEM) studies have shown that, at least a proportion of inclusions, are rich in vesicles that are coated by mostly non-fibrillar αSyn (Lashuel, 2020; Shahmoradian *et al*, 2019). The close relation between αSyn and lipids ties into recent evidence suggesting a role of phase separation in the biology of αSyn. In vitro studies have shown that αSyn forms condensates (Hardenberg *et al*, 2021; Ray *et al*, 2020). Subsequent studies in cultured cells showed that αSyn phase separation occurs in vivo with assistance from binding partners, including lipid membranes and VAMP2, a synaptic protein that interacts with αSyn to regulate synaptic vesicle organization (Agarwal *et al*, 2024; Wang *et al*, 2024).

In this manuscript, we identified mechanisms and genetic networks that are dysregulated in sporadic PD. The starting point was our recent high throughput screen through which we identified 38 genes that modulate αSyn propagation (Kara *et al*, 2021). Two of the identified genes, whose knock-down increased αSyn propagation, were of particular interest: *TAX1BP1*, an autophagy receptor involved in PINK1/Parkin mitophagy (Lazarou *et al*, 2015; Sarraf *et al*, 2022), and *ADAMTS19*, a metalloproteinase thought to be involved in the proteolysis of αSyn (Pampalakis *et al*, 2017). We show that ADAMTS19 modulates the phase separation of αSyn into inclusions, whereas TAX1BP1 controls how αSyn and lipid droplets integrate within inclusions. We also show that knock-down of *TAX1BP1* or *ADAMTS19* leads to the differential expression of numerous genes that are enriched in rare variants in patients with sporadic PD versus healthy controls. Those genes cluster within common gene expression modules that are specific to regions of the brain that develop high degrees of αSyn pathology. We propose that rare risk variants within combinations of genes result in dysfunction of entire genetic networks and associated pathways; in turn, this leads to a cascade of events resulting in neurodegeneration. Finally, we show that chloroquine, a drug whose administration in patients with rheumatoid arthritis significantly lowers their risk for development of PD (Paakinaho *et al*, 2022), dissolves αSyn inclusions and rescues the effect caused by the knock-down of *ADAMTS19* and *TAX1BP1*.

## RESULTS

### Knock-down of *ADAMTS19* and *TAX1BP1* modulates the number and size of αSyn inclusions

It is possible to recapitulate key features of αSyn inclusions in cultured cells by strategically mutating the KTKEGV repeat motifs in the protein’s sequence. Mutating three glutamic acid (E) residues into lysine (K) (E35K, E46K, E61K) enhances the interaction between αSyn and vesicles (Rovere *et al*, 2019). This results in the spontaneous formation of inclusions that entrap vesicles and have similar ultrastructure to Lewy Bodies (Shahmoradian *et al*., 2019). We used an M17D cell line expressing under doxycycline induction this triple mutant “3K” αSyn fused with Venus YFP on its C-terminus to visualize the αSyn inclusions (Dettmer *et al*, 2015; Dettmer *et al*, 2017) (fig.1B). In that cell line, αSyn inclusions first start appearing at ∼16h post-doxycycline induction and acquire their final appearance by ∼48h.

3K cells were transfected with siRNAs targeting *TAX1BP1*, *ADAMTS19* and with non-targeting (scrambled) siRNA the day after doxycycline induction. The cells were then imaged live over the course of four days, starting at 48h after doxycycline induction, at 24h intervals (fig.1C). Both knock-downs resulted in a significant increase in the number of αSyn inclusions, which was more prominent at the first timepoint but attenuated thereafter (fig. 1D). We also observed a significant decrease in the size of the inclusions, which remained consistent over the same period (fig. 1E,F). Simultaneous knock-down of *ADAMTS19* and *TAX1BP1* did not result in a significant increase in the number or decrease in the size of the inclusions in comparison to the individual knock-downs, which suggested that they belong to the same pathway (fig. 1G,H). We then quantified the amount of αSyn per cell by flow cytometry at days 1 and 2 post-transfection and found no difference in comparison to scrambled siRNA (fig. 1I). In experiments designed in the same manner, we confirmed through RNA sequencing that *TAX1BP1* and *ADAMTS19* are expressed in the 3K M17D cells at medium expression levels. We also confirmed the knock-down efficiency of the siRNAs, as well as that *TAX1BP1* and *ADAMTS19* were the top and fifth highest differentially expressed genes (DEGs) as ranked by adjusted p-value, respectively, in the corresponding knock-down experiment. A number of DEGs for each condition were also identified, as determined by adjusted p-values <0.05 (table S1).

### αSyn inclusions are partially dynamic in condensates that form and expand through phase separation and this process is modulated by knock-down of *ADAMTS19*

We observed that αSyn inclusions in the 3K M17D cell line appeared rounded and droplet-like, raising the suspicion that liquid-liquid phase separation is involved in their formation. To assess this, we undertook fluorescence recovery after photobleaching (FRAP) experiments, in which we bleached either entire inclusions or a proportion of their surface area at 48h after doxycycline induction. αSyn molecules were mobile at varying degrees within inclusions, which is consistent with them being liquid condensates formed via phase separation (fig. 2A,B).

**Figure 2:**
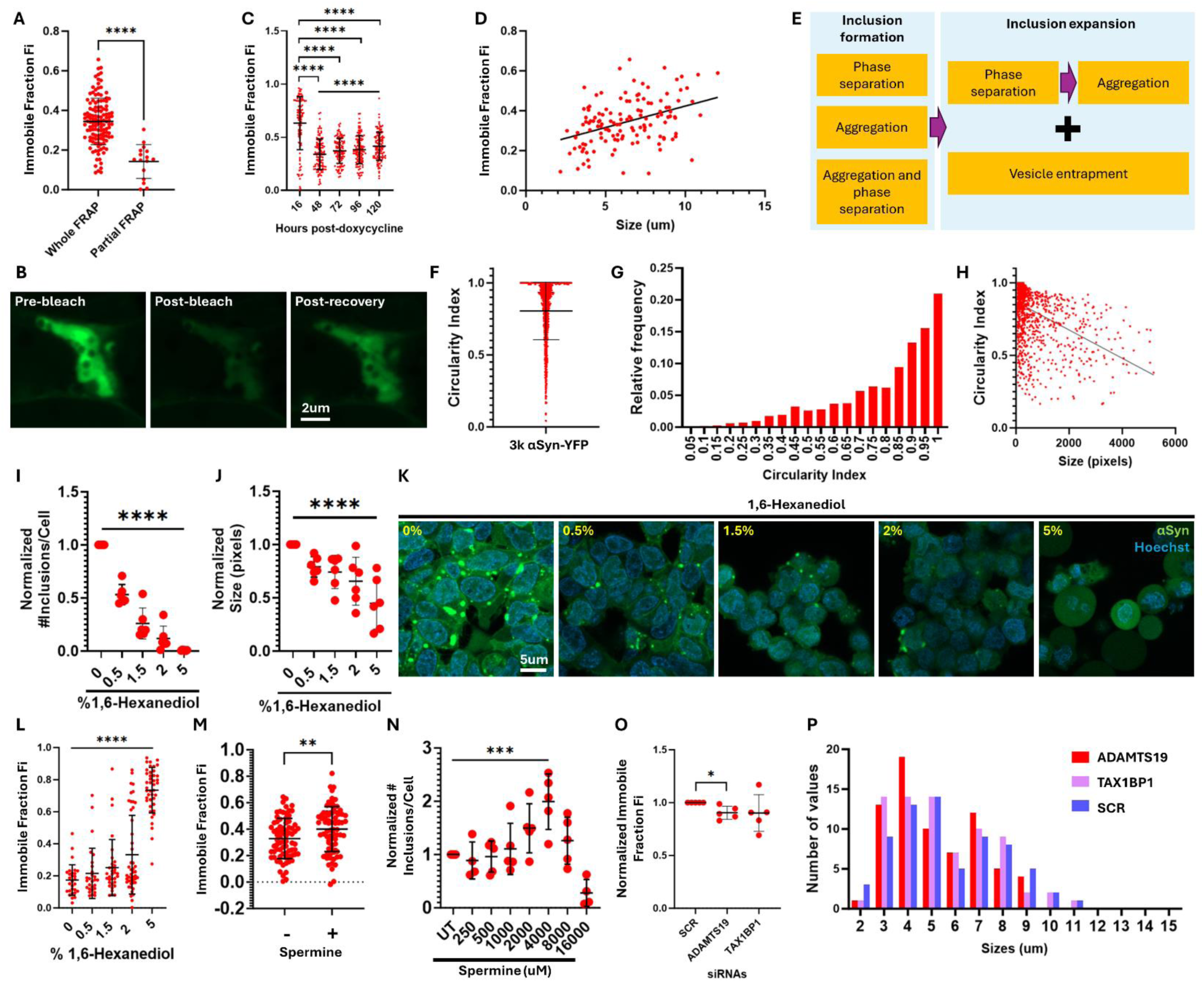
The role of phase separation in the formation of αSyn inclusions. A. Fluorescence recovery after photobleaching (FRAP) plots on the inclusions formed in the 3K αSyn M17D cell line. The inclusions were bleached either in their entirety or partially by 8 iterations of the 488nm laser and their recovery was monitored in timelapse imaging. Shown in the plot is the immobile fraction (Fi), which represents the proportion of fluorescence that did not recover, as indicated by the plateau level reached. Unpaired two tailed t-test. B. Confocal images showing the appearance of one αSyn inclusion before bleaching, right after bleaching and post-recovery (plateau phase). C. Timecourse FRAP experiment showing the evolution of the Fi over 120h. The 16h timepoint was compared to each subsequent timepoint through a one way ANOVA with Dunnett test correction for multiple testing. Timepoints 48-120h were analyzed through a one way ANOVA with test for linear trend. D. XY graph showing the relationship between the size of αSyn inclusions and their Fi. Data was analyzed through Pearson and Spearman correlation tests, both of which showed a p-value <0.0001. R squared=0.1592. Equation determined through simple linear regression: Y=0.02150*X+0.2086. E. Proposed sequence of changes in the liquid-solid status of αSyn inclusions over time. F. Plot showing the circularity index of αSyn inclusions in the 3K cell line. G. Frequency plot showing the distribution of the circularity indexes for the αSyn inclusions (bin size=0.05). H. Correlation plot for the circularity index versus inclusion size. Data was analyzed through Pearson and Spearman correlation tests, both of which showed a p-value <0.0001. R squared=0.2018. Equation determined through simple linear regression: Y=-9.74e-005*X+0.8702. I. Plot showing the number and J. size of αSyn inclusions per cell after treatment with various concentrations of 1,6-Hexanediol (in %v/v). Data was analyzed through a one way ANOVA with test for linear trend. Data was normalized to the untreated control within each independent experiment prior to putting the data together, which was arbitrarily assigned the value of 1. K. Representative confocal images of 3K cells that were treated with dose-response concentrations of 1,6-Hexanediol (in %v/v). L. FRAP experiments on αSyn inclusions after treatment with dose-response concentrations of 1,6-Hexanediol (in %v/v). Data was analyzed through a one way ANOVA with test for linear trend. M. FRAP experiments on αSyn inclusions after treatment with 4000uM of Spermine. Whole inclusions were bleached. Unpaired two tailed t-test. N. Number of αSyn inclusions per cell after treatment with dose-response concentrations of Spermine. Data was normalized to the untreated control within each independent experiment, which was arbitrarily assigned the value of 1. Data was analyzed through a one way ANOVA with test for linear trend for the concentrations between 0-4000uM of Spermine. O. FRAP experiments on αSyn inclusions after knock-down of *TAX1BP1* and *ADAMTS19* versus non-targeting scrambled control (2 days post-transfection). Data was normalized to the negative control within each independent experiment, which was arbitrarily assigned the value of 1. Data was analyzed through one sample t-tests followed by Bonferroni correction for multiple tests. P. Frequency plot showing the distribution of the sizes of the inclusions that were analyzed for each siRNA. This was done to ensure that matched populations of inclusions were imaged for each condition, given that we have observed a change in Fi depending on the size of the inclusions (see fig. 2D). Plots containing normalized data are labelled as such on the Y axis. 5 biological replicates were done for each experiment, unless stated otherwise. P values: *≤0.05, **≤0.01, ***≤0.001, ****≤0.0001. Only statistically significant differences are shown in the plots. Abbreviations: UT: untreated; SCR: scrambled; Fi: immobile fraction.

Next, we undertook a time course FRAP experiment, in which we bleached whole inclusions at the timepoint when they first became visible (16h after doxycycline induction), and subsequently at 48h, 72h, 96h and 120h. The immobile fraction (Fi) of the inclusions at 16h ranged from 0-1 and decreased significantly at subsequent timepoints. There was a significant trend for the increase of the Fi of the inclusions over time between 48-120h (fig. 2C). Finally, we observed a significant positive correlation between the size of the inclusions and their Fi (fig. 2D). These findings suggest that αSyn inclusions are initially formed either through phase separation or aggregation, or a combination thereof, but expand through phase separation. The larger αSyn inclusions are also more congealed, suggesting that when larger quantities of αSyn accumulate within inclusions it starts to aggregate. In summary, the biogenesis of αSyn inclusions follows a complex process involving intermixed steps of phase separation and aggregation (fig. 2E).

We then undertook a series of confirmatory experiments. We quantified the geometry of αSyn inclusions through the circularity index (Iannucci *et al*, 2023) and found that most of them were round (circularity index over 0.85), in keeping with a droplet nature (fig. 2F,G,H). Treatment with a dose-response of 1,6-Hexanediol concentrations, which is an aliphatic alcohol that decreases hydrophobic interactions (Zhang *et al*, 2020), showed that high concentrations were associated with reduced number and size of αSyn inclusions (fig. 2I,J,K). High concentrations of 1,6-Hexanediol were also associated with significantly higher Fi, suggesting that the liquid component is dissolved and the solid one maintained (fig. 2L). Finally, we treated cells with 4000uM of Spermine, a polyamine with four positive charges that disrupts intermolecular interactions of αSyn by binding to its negatively charged C-terminus (Agarwal *et al*., 2024). FRAP measurements showed that this treatment increases the Fi of αSyn inclusions (fig. 2M). A significant dose-response association was seen between higher Spermine concentrations and number of inclusions per cell (fig. 2N). The modulation of inclusions by 1,6-Hexanediol and Spermine, which modify intermolecular interactions mediated by hydrophobic interactions and charges, is supportive (but not diagnostic) for αSyn inclusions being liquid condensates.

Knock-down of *ADAMTS19*, but not *TAX1BP1*, resulted in a decrease of the Fi of αSyn inclusions, indicating that they become more liquid (fig. 2O,P). It is possible that, since they are more liquid, they are also more dynamic and undergo fission more readily, thereby explaining the increased number and reduced size of αSyn inclusion after *ADAMTS19* knock-down.

### αSyn condensates closely associate with lipid droplets, vesicles and membranous organelles within inclusions and this relationship is modulated by *TAX1BP1* knock-down

The brain is the second most lipid rich organ after adipose tissue, and lipid dysregulation has an important role in the pathogenesis of PD (Fanning *et al*, 2020). To study the relationship between αSyn and lipids, we simulated a lipid-rich environment in 3K cells by treating them with Oleic Acid for 16h. Treatment with Oleic Acid increases the intracellular content of neutral lipids as well as the number of lipid droplets (in which neutral lipids are sequestered) (Papadopoulos *et al*, 2015). To visualize the lipid droplets, we stained the cells with LipidTOX deep red. A proportion of αSyn inclusions in the untreated condition entrapped a small number of lipid droplets. However, treatment with Oleic Acid amplified this phenomenon and resulted in significantly higher number of lipid droplets entrapped within enlarged αSyn inclusions, imparting a “Swiss cheese” appearance (fig. 3A).

**Figure 3:**
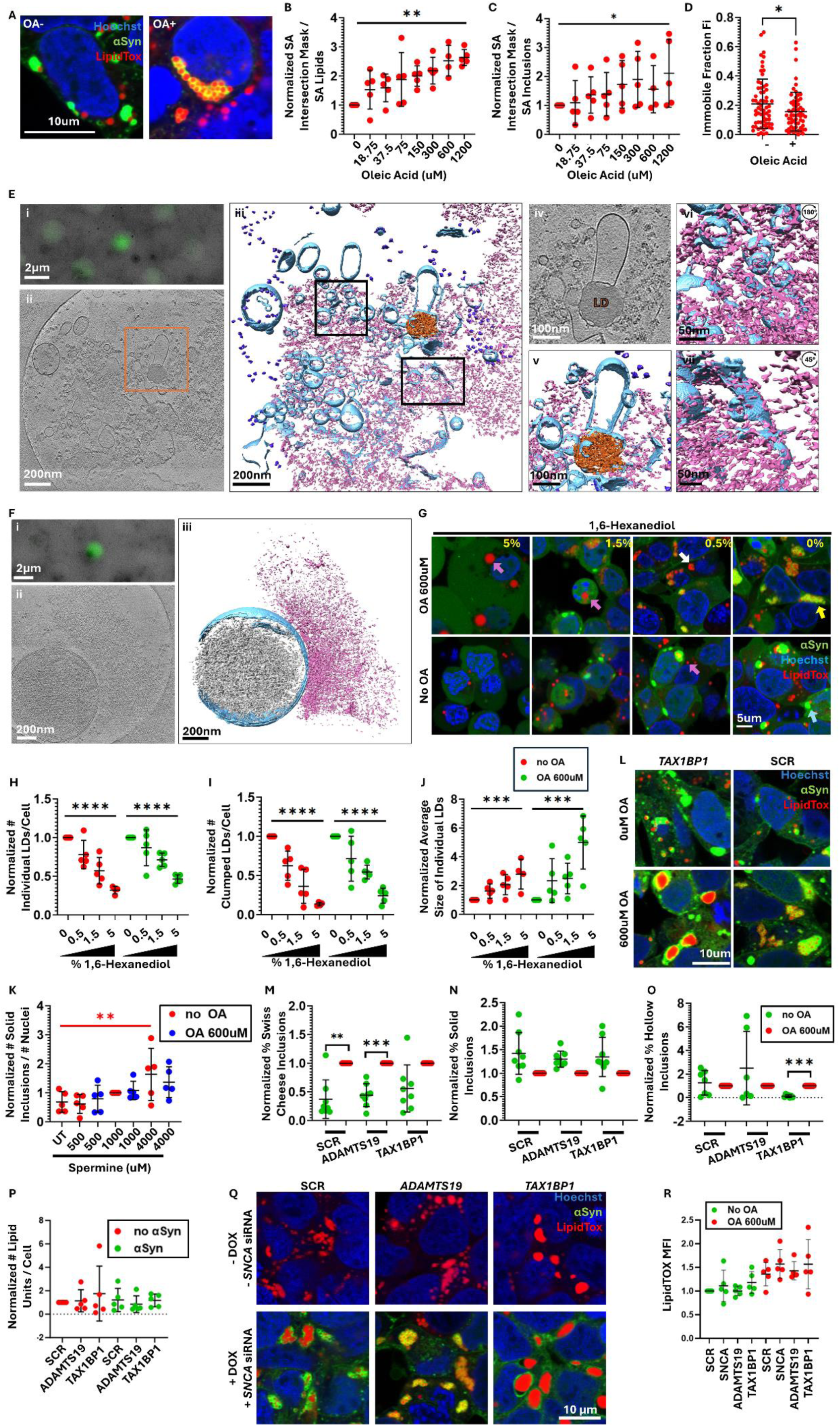
Lipid droplets, membranes and vesicles integrate within αSyn inclusions. A. Representative confocal images showing αSyn inclusions before and after treatment with 600uM of Oleic Acid (OA). Cells were stained with LipidTOX Deep Red stain for neutral lipids. B. Plot showing the fraction of the overlapping surface areas of the red (LipidTOX) and green (αSyn) masks relatively to the total surface area of the red mask. The results represent the fraction of the lipid droplets that are not embedded within αSyn inclusions. Data was analyzed through a one way ANOVA with test for linear trend. C. Plot showing the fraction of the overlapping surface areas of the green (αSyn) and red (LipidTOX) masks relatively to the total surface area of the green mask. The results represent the fraction of the αSyn inclusions that are not associated with lipid droplets. Data was analyzed through a one way ANOVA with test for linear trend. D. FRAP on lipid droplets after 16h of 600uM Oleic Acid or without Oleic Acid treatment. Lipid droplets were bleached simultaneously with 50 iterations of the 561nm and the 640nm lasers. Lipids are more mobile within lipid droplets in cells that have been treated with Oleic Acid versus untreated cells. Unpaired two tailed t-test. E. Cryo-electron tomography (cryo-ET) of cell lysates prepared from 3K M17D cells show a lipid droplet and fragmented membrane pieces meshed within amorphous αSyn aggregates. (i) Correlative light microscopy and low magnification electron microscopy (EM) image of a representative αSyn punctum. (ii) Slice view of a tomogram of the αSyn punctum in (i). (iii) Isosurface view of the tomogram. (iv) Zoom-in slice view of a lipid droplet and surrounding αSyn densities and membrane fragments as in orange box in (ii). (v) 3D Isosurface view of the lipid droplet in (iv). (vi) Isosurface view of upper left boxed in region from (iii), rotated 180° showing αSyn on the surface of membrane fragments and vesicles. (vii) Isosurface view of lower right boxed in region from (iii), rotated 45° showing αSyn densities on membranes with significant deformation. Blue: membrane; Pink: αSyn aggregate; Orange: lipid droplet; Purple: ribosomes. F. Cryo-ET of cell lysates show interactions between αSyn amorphous aggregates and mitochondrial outer membrane. (i) Correlative light microscopy and low magnification electron microscopy (EM) image of an αSyn punctum in cell lysate prepared from 3K M17D cells. (ii) Slice view of a tomogram of the αSyn punctum in (i). (iii) Isosurface view of the tomogram showing a mitochondrion with an amorphous αSyn aggregate associated with the outer membrane. Grey: mitochondria matrix; Blue: mitochondria membrane system; pink: αSyn. G. Representative confocal images of cells that were treated with dose-response concentrations of 1,6-Hexanediol (%v/v) after 16h of treatment with 600uM of Oleic Acid (OA) or no pre-treatment with OA. Yellow arrow: Swiss Cheese αSyn inclusion; Blue arrow: Solid αSyn inclusion; Purple arrows: single lipid droplet; White arrow: Clustered lipid droplets. H. Plots showing the number of individual lipid droplets per cell. For the Oleic Acid treated and untreated conditions separately, data was normalized to the untreated control within each independent experiment, which was arbitrarily assigned the value of 1. Data was analyzed through a one way ANOVA with test for linear trend. I. Plot showing the number of clustered lipid droplets per cell. For the Oleic Acid treated and untreated conditions, data was normalized to the untreated control within each independent experiment, which was arbitrarily assigned the value of 1. Data was analyzed through a one way ANOVA with test for linear trend. J. Plot showing the size of single lipid droplets. For the Oleic Acid treated and untreated conditions, data was normalized to the untreated control within each independent experiment, which was arbitrarily assigned the value of 1. Data was analyzed through a one way ANOVA with test for linear trend. K. Plot showing the normalized number of solid αSyn inclusions divided by the number of nuclei. Data was normalized to the 1000uM Spermine condition within each independent experiment, which was arbitrarily assigned the value of 1. OA-treated and untreated cells were analyzed separately through one way ANOVA with one way test for trend. This was significant only for the non OA treated condition (shown in red). L. Representative confocal images showing the effect of *TAX1BP1* or *ADAMTS19* knock-down on the lipid droplets in the presence or absence of αSyn protein. M. Plot showing the normalized proportion of Swiss cheese inclusions after knocking down *TAX1BP1* or *ADAMTS19*, versus scrambled control, in the presence or absence of Oleic Acid treatment. Data was normalized to the untreated sample for each siRNA, which was designated arbitrarily as 1. Data was analyzed through one sample t-tests followed by Bonferroni correction for multiple tests. 8 biological replicates. N. Plot showing the normalized proportion of solid inclusions after knocking down *TAX1BP1* or *ADAMTS19*, versus scrambled control, in the presence or absence of Oleic Acid treatment. Data was normalized to the untreated sample for each siRNA, which was designated arbitrarily as 1. Data was analyzed through one sample t-tests followed by Bonferroni correction for multiple tests. 8 biological replicates. O. Plot showing the normalized proportion of hollow inclusions after knocking down *TAX1BP1* or *ADAMTS19*, versus scrambled control, in the presence or absence of Oleic Acid treatment. Data was normalized to the untreated sample for each siRNA, which was designated arbitrarily as 1. Data was analyzed through one sample t-tests followed by Bonferroni correction for multiple tests. 8 biological replicates. P. Number of lipid units (including single and clumped lipid droplets) per cell after knocking down *TAX1BP1* or *ADAMTS19* versus scrambled control, in the presence and absence of endogenous αSyn. Data was normalized to the SCR siRNA, which was designated arbitrarily as 1. Samples in the presence and absence of endogenous αSyn were analyzed separately through a one way ANOVA with Dunnett test correction for multiple testing. Q. Representative confocal images for (P). R. Flow cytometry experiments showed that knock down of *TAX1BP1* or *ADAMTS19* did not affect the amount of intracellular lipids (as indicated by the MFI of LipidTOX neutral lipid dye) 2 days after transfection, either with or without treatment with 600uM of Oleic Acid. One way ANOVA with Dunnett test correction for multiple testing. Data was normalized to the untreated control within each independent experiment, which was arbitrarily assigned the value of 1, unless stated otherwise. Plots containing normalized data are labelled as such on the Y axis. 5 biological replicates were done for each experiment, unless otherwise stated. P-values: *≤0.05, **≤0.01, ***≤0.001, ****≤0.0001. Only statistically significant differences are shown in the plots. Abbreviations: OA: Oleic Acid; SA: surface area; LDs: lipid droplets; DOX: doxycycline; MFI: mean fluorescence intensity.

Analysis of co-localization between the green (αSyn) and red (LipidTOX) channels showed that the proportion of αSyn and lipid droplets that associate with each other significantly increases as the concentration of Oleic Acid increases (fig. 3B,C). This suggests that the free fraction of lipid droplets and αSyn decreases when the intracellular contents of neutral lipids increase; in other words, lipid droplets preferentially associate with αSyn inclusions and αSyn preferentially associates with lipid droplets. FRAP measurements on LipidTOX-positive droplets showed that the neutral lipids are mobile within droplets, which is consistent with prior findings that they form through phase separation upon budding off of the endoplasmic reticulum (fig. 3D) (Zoni *et al*, 2021). Correlative light and electron microscopy (CLEM) followed by cryo-ET on αSyn inclusions that were isolated by hypotonic lysis buffer showed vesicles, lipid droplets and membranous organelles, such as mitochondria, that are coated by amorphously aggregated αSyn (fig. 3E,F), consistent with previous reports on the 3K cell line (Dettmer *et al*., 2015; Dettmer *et al*., 2017). Membrane components entrapped within the amorphous network of alpha-Syn aggregates exhibited marked deformation and disruption of membrane integrity. This disruption corresponds to findings that higher-order structures formed by intrinsically disordered proteins can directly interact with and penetrate membranes, impairing the functions of cellular organelles, such as mitochondria, autophagosomes, and ER (Bauerlein *et al*, 2017). The close association between αSyn condensates and the surface of lipid droplets/vesicles/membranous organelles within inclusions suggests that the phase separation of αSyn could be initiated by the binding between αSyn and lipids. This is in keeping with a recent report that the formation of VAMP2-mediated αSyn condensates is dependent on the binding between αSyn and lipids (Agarwal *et al*., 2024).

1,6-Hexanediol showed a significant dose-response effect on the number of isolated, single lipid droplets and on the number of clusters of lipid droplets, with higher concentrations being associated with reduction in number (fig. 3G,H,I). A significant dose-response effect was also seen on the size of the isolated single lipid droplets (fig. 3J) but not on the size of lipid droplet clusters (fig. S1A). Finally, a significant dose-response effect was seen on the overlap between αSyn inclusions and lipid droplets, with higher concentrations reducing the overlap (fig. S1B,C). Spermine treatment showed a significant dose-response effect on the total number of inclusions per cell (fig. S1D) and on the number of solid inclusions per cell in the absence of Oleic Acid treatment, with higher concentrations associated with higher number of inclusions. However, this effect disappeared after Oleic Acid treatment (fig. 3K). The fact that lipid droplets and their association with αSyn inclusions is modulated by 1,6-hexanediol and spermine treatment supports a role for phase separation in those behaviors.

Next, we set out to determine whether knock-down of *ADAMTS19* or *TAX1BP1* modifies how lipids integrate within αSyn inclusions. Knock-down of *TAX1BP1* resulted in large lipid droplets surrounded by a thin rim of αSyn (“hollow” inclusions) (fig. 3L,M,N,O). However, in the complete absence of αSyn, knock-down of *TAX1BP1* and *ADAMTS19* had no effect on lipid droplets (fig. 3P,Q). Those findings suggest that TAX1BP1 initially impacts on αSyn, which, in turn, modulates the lipid droplets. Finally, experiments on iPSc-dopaminergic neurons from patients with *SNCA* triplication versus isogenic controls showed no difference between the number of lipid droplets per cell, irrespectively of Oleic Acid treatment status, suggesting that the intracellular amount of αSyn within neurons does not have an impact on lipid droplets (fig. S1E,F,G,H). Of note, flow cytometry experiments showed that the amount of neutral lipids per cell does not change when *TAX1BP1* is knocked down, which suggests that the total quantity of neutral lipids does not have a role in the changes mediated by *TAX1BP1* knock-down (fig. 3R).

### The mutations in the 3K αSyn protein increase its propensity to phase separate and associate with phospholipids

While 3K αSyn is an artificial mutant, it does result in spontaneous formation of inclusions that entrap vesicles and recapitulate ultrastructural features of Lewy Bodies (Shahmoradian *et al*., 2019). We undertook experiments and analyses to understand the molecular grammar of αSyn and how the 3K mutant modulates interaction with lipids and phase separation. This knowledge is useful to draw comparisons between the 3K model and the in vivo situation in humans and assist with the interpretation of experimental findings.

In vitro experiments confirmed the increased propensity of the 3K αSyn protein to phase separate in comparison to the wild type protein, as it formed condensates at lower concentrations of crowding agent (∼7% lower PEG concentration) (fig. 4A). We next investigated whether this is caused by the increased content in lysine (Ukmar-Godec *et al*, 2019) or by the disruption of the KTKEGV repeat motifs. We cloned a construct encoding αSyn with intact KTKEGV repeat motifs but with one extra lysine right before the 3^rd^,4^th^ and 5^th^ repeat motifs (KKTK αSyn), which has the same lysine content as the 3K protein (fig. 4B). Transfection of M17D cells, with and without subsequent treatment with Oleic Acid, showed that the KKTK protein forms neither inclusions nor conglomerates with lipids (fig. 4C,D). This indicates that disruption of the KTKEGV repeat motifs is the initiating event for the phase separation for the 3K αSyn.

**Figure 4:**
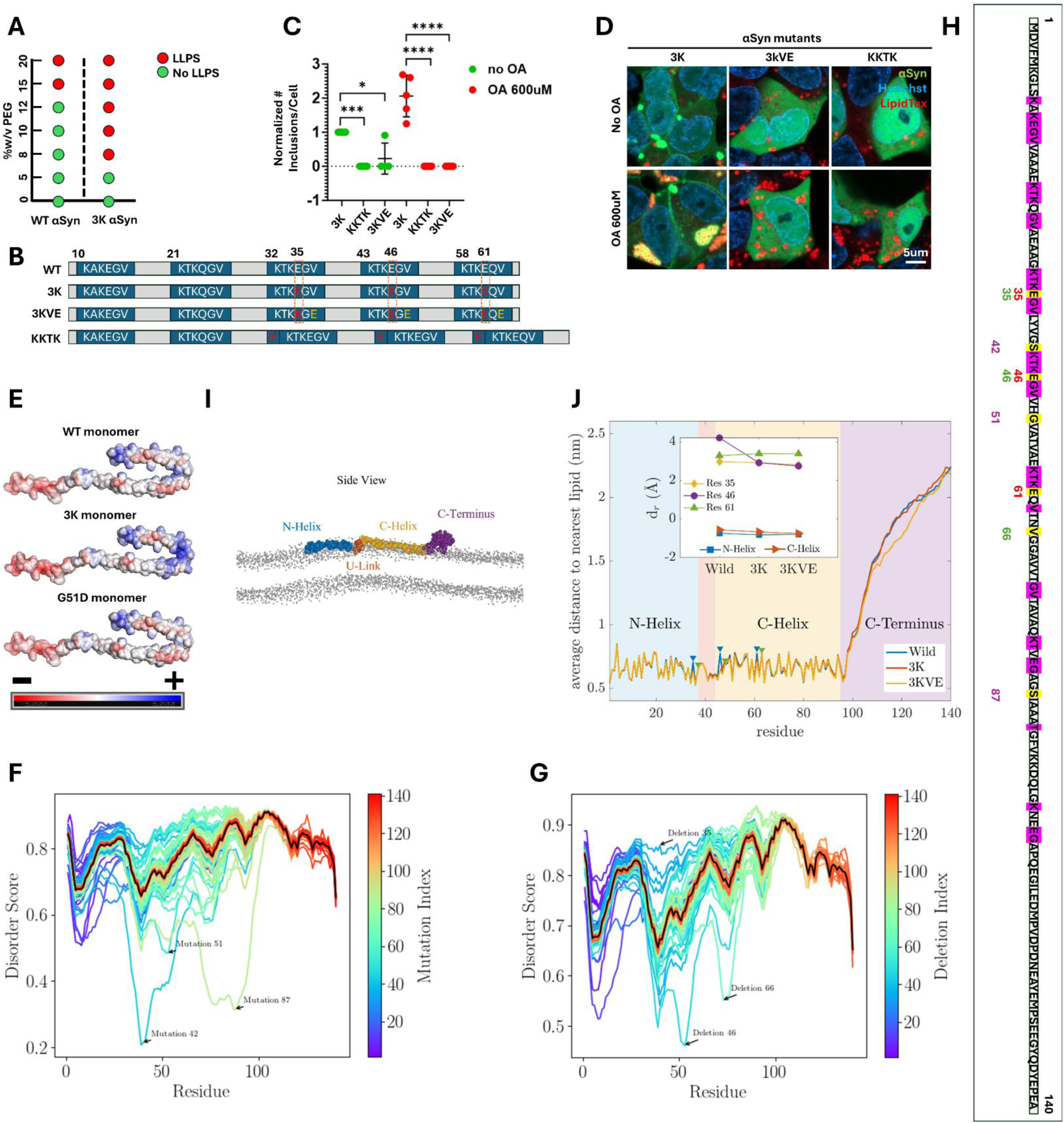
The propensity of 3K αSyn to phase separate and interact with lipids. A. Phase separation diagram showing the phase separation of αSyn at various concentrations of crowding agent PEG-8000 and αSyn protein (wild type versus 3K mutant). B. Diagrams of the amino acid sequence of αSyn showing the wild type (WT) version, as well as the artificial mutants studied. The repeat motifs are in blue rectangles. In red are the glutamic acid to lysine substitutions. In yellow are the valine to glutamic acid substitutions. C. Number of αSyn inclusions per cell for the three artificial mutants: 3K, KKTK, 3KVE. Data was normalized to the 3K sample without Oleic Acid treatment, which was arbitrarily designated as 1. Data was analyzed through a one way ANOVA with Dunnett test correction for multiple testing, separately for the OA-treated and untreated conditions. P-values: *≤0.05, **≤0.01, ***≤0.001, ****≤0.0001. Only statistically significant differences are indicated on the plot. D. Representative confocal images of the 3K, 3KVE and KKTK αSyn mutants, with and without Oleic Acid treatment. E. Diagrams showing the electrostatic charges for monomeric αSyn (wild type, 3K and G51D mutants). F. Plot showing the disorder score for the αSyn protein after sequentially mutating one by one every residue into an amino acid with opposite properties. G. Plot showing the disorder score for the αSyn protein after sequentially deleting one by one every residue. H. Diagram of the amino acid sequence of the αSyn protein showing the artificial mutations studied. Highlighted in pink are the imperfect KTKEGV repeat motifs and in yellow the mutations. In red font are the mutated residues in the 3K protein, in green and in purple are the residues whose deletion or substitution with amino acids with opposite properties, respectively, was found to profoundly impact disorder in silico. I. Side view of coarse-grained representation of the wild-type αSyn in the Martini model. Different regions of the protein including the N-Helix (residues 1-33), U-Link (residues 34-44), C-Helix (residues 45-92), and the C-Terminus (residues 93-140) are shown in blue, orange, yellow, and purple, respectively. The head groups of POPG lipids are shown in grey. J. The average distance of each residue to the nearest lipid during the 6μs-long simulation. Blue triangle markers correspond to the mutated residues 35, 46, 61, and green markers indicate the mutated residues 37, 48, 63, both with respect to the wild type alpha-synuclein. The inset shows the relative depth d_r_ of the mutated residues and the centres of mass of the N- and C-Helices with respect to the lipid bilayer’s top surface for each mutant. Abbreviations: WT: Wild type; LLPS: liquid liquid phase separation; OA: Oleic Acid; POPG: palmitoyloleoylphosphatidylglycerol.

We then focused on the molecular mechanism with which 3K αSyn interacts with lipids. We first plotted the electrostatic charges of monomeric wild type and 3K αSyn (fig. 4E). There was a cluster of positively charged residues located around residue 43 in the hairpin fold in the monomer, which are more positively charged in the 3K and more negatively charged in the G51D mutant (Kiely *et al*, 2013) compared to the wild type protein, respectively. We hypothesized that the Valine (V) residues within the KTKEGV repeat motifs are crucial for the interaction with lipids, given their hydrophobicity. We substituted the hydrophobic Valine residues 37, 48 and 63 in the 3^rd^, 4^th^ and 5^th^ repeat motifs with negatively charged glutamic acid (E), which is expected to repel lipids (3KVE αSyn) (fig. 4B). This mutant αSyn was unable to form inclusions and integrate with lipids in conglomerates, indicating that the Valine residue is critical for both those functions (fig. 4C,D). Finally, we plotted the frequency of the various amino acids in each region of the wild type αSyn protein. The most frequent amino acid in the amphipathic region was lysine (K) (positively charged) (18%), in the NAC domain valine (V) (hydrophobic) (23.5%) and in the acidic tail glutamic acid (E) (negatively charged) (22.2%) (fig. S2A). The fact that the amphipathic region and the NAC domains are enriched in positively charged and hydrophobic residues is consistent with previous reports indicating that they are the critical regions of the protein for the interactions with lipids. It is likely that this is enhanced in the 3K αSyn due to the extra positive charges. Those findings underscore the importance of positively charged lysines, hydrophobic valines and the imperfect KTKEGV repeat motifs for the interaction between αSyn and lipids.

We augmented our approach with Molecular Dynamic simulations. We simulated the interaction between a single αSyn protein (wild type, 3K) and bilayer membranes composed of anionic palmitoyloleoylphosphatidylglycerol (POPG) lipids (fig. 4I). Those analyses further highlighted the importance of the extra lysines in the 3K αSyn protein in terms of modulating the protein’s conformation and interaction with lipids. The lysine at residue 46 in 3K αSyn has an increased distance (∼3 Angstroms) from the N-Helix (residues 1-33) in comparison to the wild type protein. We next quantified the number of lipids that are in contact with each residue and found that the E35K and E46K point mutations in the 3K mutant show an increased contact with lipids. The accumulated contact maps also showed that the E35K, E46K and E61K mutants in the 3K protein are more deeply inserted into the lipid bilayer in comparison to the wild type αSyn, with the most profound difference observed for residue 46 (fig. 4J). These results confirm the importance of the lysines at residues 35, 46 and 61 for the interaction between αSyn and lipids. See supplementary text file for further details.

We used frequency plots to plot the number of single nucleotide polymorphisms (SNPs) normalized to the length of the genomic sequence in exons, introns and the entire gene for every gene in the genome as catalogued in Ensembl genome browser. The *SNCA* gene showed lower than average number of SNPs within exons and introns (fig. S2B). This suggests that αSyn tolerates less genetic variability than most genes and that it follows a strict molecular grammar. We made the data on the normalized number of SNPs for every gene in the genome publicly available in a searchable browser *<u>karalab.shinyapps.io/SNP_Browser/</u>*. Finally, we generated *in silico* αSyn mutants in which each amino acid was sequentially either deleted or mutated into an amino acid with opposite properties (charge, polarity, hydrophobicity) (table S2) and plotted the disorder score as calculated by Metapredict (Emenecker *et al*, 2021). We found that alterations in the region of the protein between residues 0 and 100 had the most profound effect on its disorder, with hotspots in residues 35, 46 and 66 for deletions and residues 42, 51 and 87 for substitutions (fig. 4F,G,H). Of note, those residues are located with the region of αSyn containing the repeat motifs, further underscoring their importance for the biophysical behavior of the protein. Deletions of either one by one or various combinations of the first five repeat motifs similarly impacted on the disorder of αSyn (fig. S2C).

### Mitochondrial health and energy production is affected by *TAX1BP1* and *ADAMTS19* knock-down

Dysregulation of mitochondrial bioenergetics has been reported in several neurodegenerative diseases, and in particular PD (Choi *et al*, 2022; Kara *et al*., 2021; Ludtmann *et al*, 2016; Ludtmann *et al*, 2018). As a starting point to assess mitochondrial health, we measured the mitochondrial membrane potential (ΔΨ_m_) in 3K M17D cells that were not induced with doxycycline. We loaded the cells with tetramethylrhodamine methyl ester (TMRM) for 40min and then acquired z-stacks on live cells. We found that *TAX1BP1* knock-down resulted in an increase in ΔΨ_m_ in comparison to the scrambled control (fig. 5A,B). To examine the mechanism with which ΔΨ_m_ is maintained, we carried out a time course experiment in which we first treated the cells with oligomycin, which is a complex V inhibitor, followed by treatment with rotenone, a complex I inhibitor, and Carbonyl cyanide p-trifluoromethoxyphenylhydrazone (FCCP), a mitochondrial uncoupler. The pattern of the decay curves of TMRM fluorescence was the same between the knock-downs and the negative control (fig. 5C).

**Figure 5:**
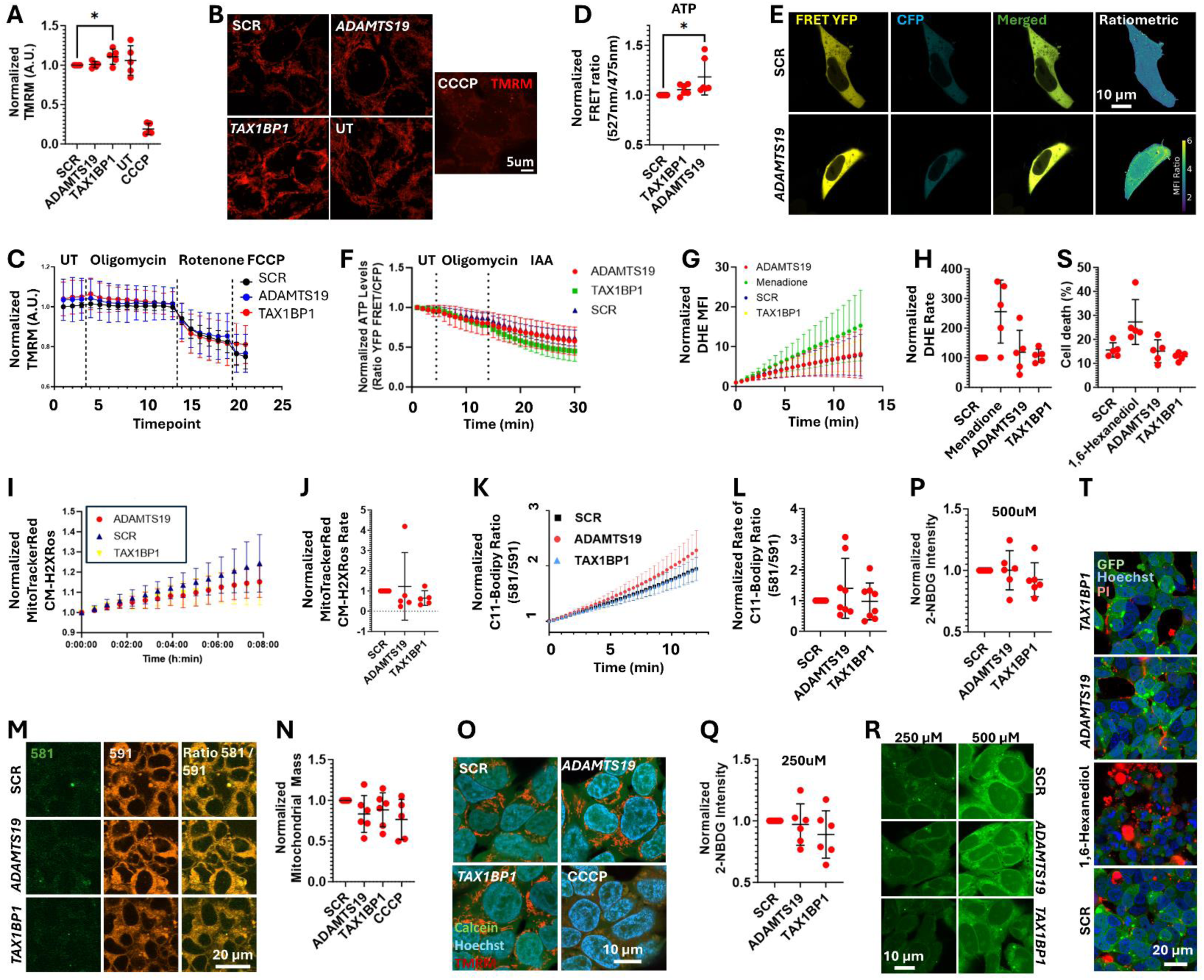
Mitochondrial bioenergetics are impacted by TAX1BP1 or ADAMTS19 knock down. A. Plot showing the mean fluorescence intensity (MFI) of TMRM (tetramethylrhodamine methyl ester) after knock-down of *TAX1BP1* or *ADAMTS19*, versus scrambled (SCR) control. CCCP (carbonyl cyanide m-chlorophenyl hydrazone) was used as a positive control, and showed a statistically significant reduction in TMRM MFI in comparison to the untreated control. Z-stacks were acquired and the maximum intensity projection images quantified. Data was analyzed through a one way ANOVA with Dunnett test correction for multiple testing, separately for the OA-treated and untreated conditions; only the SCR, *ADAMTS19* and *TAX1BP1* samples were included. B. Representative confocal images for the data plotted in (A). C. Timecourse experiment measuring the MFI of TMRM after treatment with oligomycin, rotenone and FCCP. Single plane images were taken for each timepoint. D. FRET (Forster Resonance Energy Transfer) of the ATP FRET reporter used to quantify the amount of intracellular ATP. E. Representative confocal images of the ATP FRET reporter and FRET channel from (D). Ratiometric image of FRET channel normalized to CFP channel is also shown. F. Timecourse ATP FRET measurements for each of the knockdowns versus scrambled control after treatment with oligomycin and iodoacetic acid (IAA). The timepoints for each siRNA were internally normalized to the first timepoint, which was arbitrarily designated as 1. To facilitate illustration of any differences between the slopes for each sample, it was easier for each curve to start at the same value on the Y axis. G. Timecourse DHE (dihydroethidium) MFI measurements for each of the knockdowns versus scrambled control and positive control (menadione). Timepoints were normalized to the first timepoint within each sample separately, as described in (F). H. Plot showing the slopes for each condition analyzed in (G). I. Timecourse of the MitoTrackerRed CM-H2XRos MFI for each of the siRNAs studied. Timepoints were normalized to the first timepoint within each sample separately, as described in (F). J. Plot showing the slopes for each condition analyzed in (I). K. Timecourse of the C11-Bodipy 581/591 MFI ratio for each of the siRNAs studied. Timepoints were normalized to the first timepoint within each sample separately, as described in (F). L. Plot showing the slopes for each condition analyzed in (K). M. Representative confocal images for (K,L). N. Mitochondrial mass as assessed through the volumetric ratio of TMRM/calcein through z-stack experiments. O. Representative confocal images for (N). P. 2-NBDG (2-(N-(7-Nitrobenz-2-oxa-1,3-diazol-4-yl)Amino)-2-Deoxyglucose) MFI for 500uM and Q. 250uM of the fluorescent glucose analogue to assess the impact of knocking-down the genes of interest on the efficiency of the glucose uptake. R. Representative confocal images for (P,Q). S. Plot showing the percentage of dead cells after transfection of each siRNA. 5%v/v of 1,6-Hexanediol was used as a positive control. Dead cells take up propidium iodide (PI) dye, but alive ones do not. T. Representative confocal images for (S). Data was normalized to the scrambled (SCR) control within each independent experiment, which was arbitrarily assigned the value of 1, unless stated otherwise in the legend. Plots containing normalized data are labelled as such on the Y axis. 5 biological replicates were done for each experiment, unless otherwise stated. P-values: *≤0.05, **≤0.01, ***≤0.001, ****≤0.0001. Only statistically significant differences are shown in the plots. Abbreviations: MFI: Mean fluorescence intensity; TMRM: tetramethylrhodamine methyl ester; CCCP: carbonyl cyanide m-chlorophenyl hydrazone; UT: untreated; SCR: scrambled; FRET: Forster Resonance Energy Transfer; A.U.: arbitrary units; FCCP: Carbonyl cyanide p-trifluoromethoxyphenylhydrazone; IAA: iodoacetic acid; DHE: dihydroethidium; PI: propidium iodide; 2-NBDG: 2-(N-(7-Nitrobenz-2-oxa-1,3-diazol-4-yl)Amino)-2-Deoxyglucose.

Mitochondrial respiration is associated with energy production through oxidative phosphorylation. Therefore, impairment of mitochondrial health could result in altered levels of intracellular ATP. We measured the intracellular amounts of ATP using a FRET-based ATP indicator (Kotera *et al*, 2010). Even though *TAX1BP1* knock-down cells showed alterations in mitochondrial membrane potential, their intracellular ATP content did not change. However, *ADAMTS19* knock-down resulted in increased ATP levels (fig. 5D,E). In a time course experiment, we treated cells consecutively with oligomycin and iodoacetic acid (IAA), which inhibits glycolysis, to assess the relative contribution of oxidative phosphorylation and glycolysis to ATP production through the decay curve in ATP levels. We did not observe a difference in comparison to the scrambled negative control (fig. 5F).

Reactive oxygen species (ROS) are mainly produced by mitochondria and are a source of cellular toxicity. Therefore, an increase in ROS could be a consequence of poor mitochondrial health. We measured the rate of ROS production in the cytoplasm by using dihydroethidium (DHE), a dye that detects superoxide (fig. 5G,H), and in the mitochondria by using MitoTracker Red CM-H2XRos, a mitochondrial dye that becomes fluorescent when oxidized (fig. 5I,J). We also measured the rate of lipid peroxidation by using a ratiometric Bodipy 581/ 591 C11 dye (fig. 5K,L,M). We did not observe a difference between the knock-down conditions and the negative control in any of those experiments.

Mitochondrial mass was measured through z-stacks by determining the ratio of the volume of the mitochondrial network to the total volume of the cytoplasm (indicated through calcein stain) excluding the nucleus. There was no difference between the knock-downs and the negative control (fig. 5N,O). Therefore, a relative increase in mitochondrial volume does not account for the increase in ATP production seen in the *ADAMTS19* knock-down. Glucose uptake, as assessed through the fluorescent glucose analogue 2-(N-(7-Nitrobenz-2-oxa-1,3-diazol-4-yl)Amino)-2-Deoxyglucose (2-NBDG) was also not altered in the knock-down conditions (fig. 5P,Q,R). To determine whether knock-down of *ADAMTS19* and *TAX1BP1* resulted in increased cell death, we quantified cellular viability through the propidium iodide stain and calculated the proportion of cells that have taken up the dye and, therefore, are dead or in the process of dying. Again, there was no difference compared to the negative control (fig. 5S,T).

### Chloroquine modifies the number and immobile fraction of αSyn inclusions

Chloroquine is an antimalarial drug and a disease-modifying anti-rheumatic drug (DMARD). An epidemiological study has found that patients with rheumatoid arthritis who receive chloroquine treatment have a significantly lower risk for the development of PD (Paakinaho *et al*., 2022). It is thought that people who eventually develop clinical PD exhibit neuropathological changes up to 20 years before disease onset (Savica *et al*, 2010). We hypothesized that chloroquine administration would rescue αSyn dyshomeostasis in the context of the 3K cell line model system. Accordingly, we treated 3K M17D cells with 50uM chloroquine for 3-5h at 48h after doxycycline induction, which is the timepoint at which αSyn inclusions are well formed. We observed a significant decrease in the number of αSyn inclusions per cell, including after knocking down *TAX1BP1* or *ADAMTS19* (fig. 6A,C,E).

**Figure 6:**
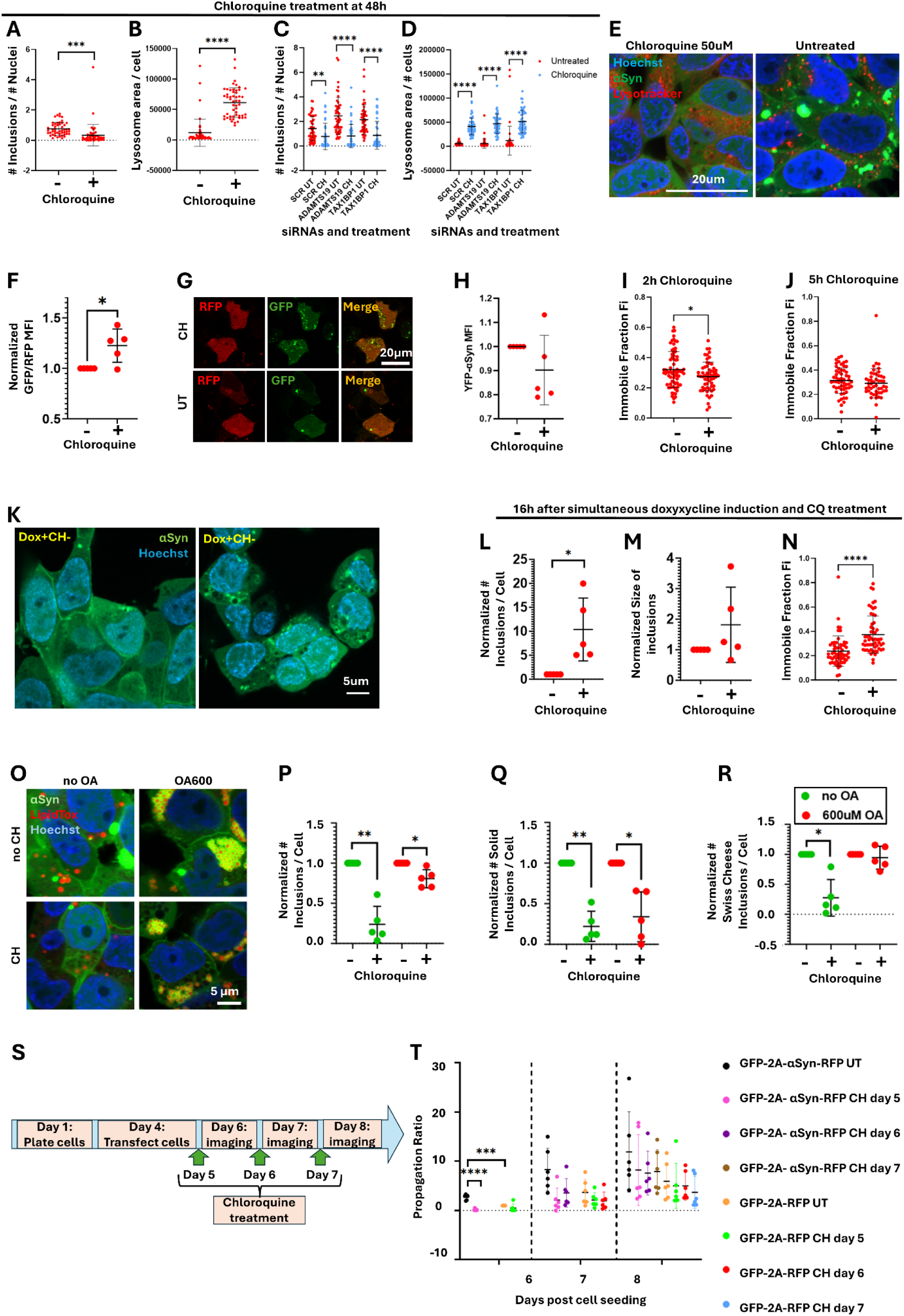
Effect of Chloroquine treatment on αSyn inclusions. A. Cells were treated with 50uM of chloroquine for 2h at 48h after doxycycline induction and the number of αSyn inclusions per cell B. And lysosomal surface area was quantified. Unpaired two tailed t-tests. C. Number of αSyn inclusions per cell D. and lysosomal surface area with and without treatment with 50uM of chloroquine at 48h after doxycycline induction and siRNA knock-down of *ADAMTS19* and *TAX1BP1*. Unpaired two tailed t-tests. E. Representative confocal images for (A,B). F. Uninduced M17D cells were transfected with a construct encoding GFP-LC3-RFP-LC3ΔG; the protein product is cleaved by endogenous ATG4 proteases into GFP-LC3 and RFP-LC3ΔG. GFP-LC3 is digested through autophagy, whereas RFP-LC3ΔG remains in the cytoplasm. The GFP/RFP ratio is a measure of autophagy activation, with a lower ratio representing a higher autophagy rate. Chloroquine treatment resulted in an increased GFP/RFP ratio in comparison to untreated control, indicating a blockage in autophagy. One sample t-test. G. Representative confocal images of (F). H. Flow cytometry experiments showed that the MFI of YFP did not change after 5h of chloroquine treatment, indicating that chloroquine treatment does not affect the amount of αSyn per cell. One sample t test. I. Effect of 2h chloroquine treatment at 48h after doxycycline induction on the immobile fraction of αSyn inclusions, as determined by FRAP experiments. Unpaired two tailed t-test. J. Effect of 5h chloroquine treatment at 48h after doxycycline induction on the immobile fraction of αSyn inclusions, as determined by FRAP experiments. Unpaired two tailed t-test. K. Representative confocal images showing αSyn inclusions that formed 16h after simultaneous doxycycline induction and chloroquine treatment. L. Quantification of (K) showing that chloroquine treated cells showed a larger number of αSyn inclusions per cell, M. but there was no difference in the average size of the inclusions. N. FRAP measurements showed that chloroquine treatment significantly increased the immobile fraction of the inclusions. (L-M) were analyzed through one sample t-tests and (N) by unpaired two tailed t-test. O. Representative confocal images of 3K cells after 16h of 600uM Oleic Acid treatment and 5h of 50uM chloroquine treatment. All four combinations of treatment statuses were tested. P. Plot showing the total number of αSyn inclusions per cell, Q. the number of solid αSyn inclusions per cell and R. the number of Swiss cheese αSyn inclusions per cell after 5h of chloroquine treatment in comparison to the untreated condition, and in the presence or absence of pre-treatment with 600uM of Oleic Acid. Samples that were and were not pre-treated with Oleic Acid were normalized separately to the non-chloroquine treated condition, which was arbitrarily designated as 1. One sample t-tests. S. Diagram showing the workflow of the chloroquine experiment in cultured cells that were transfected with the GFP-2A-αSyn-RFP or the GFP-2A-RFP constructs to monitor αSyn propagation. Cells were seeded in coverglass bottom chamberslides on day 1. T. Propagation ratio of αSyn-RFP and RFP alone. To determine the propagation ratio, the green mask was subtracted from the red mask to identify the regions that contained only RFP (and thereby propagating protein); the surface area of those regions was divided with the surface area of the green mask for within-image normalization. Prior to putting all the data from independent experiments together, data was normalized to the GFP-2A-RFP negative control at day 6, which was arbitrarily designated as 1. For an explanation on what the number of days correspond to, please refer to panel S. 6 biological replicates. Two way ANOVA with Tukey’s correction. Data was normalized to the untreated control within each independent experiment, which was arbitrarily assigned the value of 1, unless stated otherwise. 5 biological replicates were done for each tissue culture experiment, unless otherwise stated. Plots containing normalized data are labelled as such on the Y axis. P-values: *≤0.05, **≤0.01, ***≤0.001, ****≤0.0001. Only statistically significant differences are shown in the plots. Abbreviations: WT: wild type; CQ: chloroquine; FRAP: fluorescence recovery after photobleaching; OA: Oleic Acid; UT: untreated.

We investigated autophagy modulation as a mechanism with which chloroquine treatment could impact on αSyn inclusions. Chloroquine inhibits autophagy by blocking the fusion between autophagosomes and lysosomes (Mauthe *et al*, 2018) and increasing the intraluminal pH in lysosomes (Fedele & Proud, 2020). Transfection with a construct encoding GFP-LC3-RFP-LC3ΔG to monitor autophagy activation (Kaizuka *et al*, 2016) showed an increased GFP/RFP ratio in chloroquine treated cells in comparison to untreated control, indicating a blockage in autophagy, as expected (fig. 6F,G). Staining with Lysotracker showed that the surface area of the lysosomes is significantly increased in cells that were treated with chloroquine (fig. 6B,D,E). This is likely caused by the accumulation of undigested material due to the block in autophagy (Fedele & Proud, 2020). Additionally, quantification of the amount of αSyn per cell through flow cytometry showed no significant reduction after chloroquine treatment (fig. 6H). These results indicate that autophagy modulation through chloroquine treatment likely does not have a role in clearing αSyn inclusions and that there is an alternative mechanism in play.

Given that αSyn inclusions in the 3K M17D cell line form and expand through phase separation, we hypothesized that chloroquine modulates the phase separation behavior of αSyn as a mechanism for αSyn inclusion dissolution and removal. We undertook FRAP measurements on αSyn inclusions before and after 2h and 5h treatment with 50uM of chloroquine. Chloroquine significantly decreased the Fi of the inclusions after treatment for 2h, indicating that they become more liquid and raising the possibility that chloroquine dissolves them (fig. 6I,J). We then assessed whether the timing of chloroquine treatment had a differential impact on αSyn inclusions. We added chloroquine simultaneously with doxycycline induction in 3K cells and imaged them 16h later, which is the timepoint at which αSyn inclusions first become visible. We observed a significant increase in the number of αSyn inclusions but no change in their size (fig. 6 K,L,M). FRAP measurements showed that chloroquine treatment significantly increased the Fi of those inclusions (fig. 6N). Those effects were opposite to what was seen for chloroquine treatment at 48h post-doxycycline induction.

Given the lipid-rich nature of the human brain and of Lewy Bodies (Shahmoradian *et al*., 2019), we assessed whether lipid droplet-rich inclusions can be cleared efficiently by chloroquine treatment. We treated 3K cells with 600uM of Oleic Acid for 16h, followed by 5h with chloroquine. Oleic Acid treatment prevented the removal of Swiss cheese αSyn inclusions, but not of the solid inclusions (fig. 6. O,P,Q,R).

We also assessed whether chloroquine modulates αSyn propagation. We transfected HEK cells with our pcDNA3.1-eGFP-2A-αSyn-RFP propagation reporter and the negative control pcDNA3.1-eGFP-2A-RFP (Kara *et al*., 2021). One day later, we treated the cells with 50uM of chloroquine and imaged them over the following three days at 24h intervals. We also treated cells with chloroquine at 48h and 72h after transfection to assess the impact of chloroquine treatment at various stages of αSyn propagation (fig. 6S). There was a significant reduction of αSyn propagation when chloroquine treatment was added one day after transfection and propagation was measured one day later. For the other conditions, there was a non-significant trend for reduction in propagation. The cell-to-cell transfer of RFP alone (negative control) was not modified by chloroquine treatment (fig. 6T).

### *TAX1BP1* and *ADAMTS19* integrate within genetic networks carrying an increased burden of rare variants in patients with PD

To get a deeper insight into the mechanisms of action of *TAX1BP1* and *ADAMTS19*, we used Weighted Gene Co-expression Analyses (WGCNA) (Langfelder & Horvath, 2008) on gene expression data from healthy human brains. This technique groups genes in modules based on their expression patterns (Konopka *et al*, 2012). It is useful in identifying genes that are co-expressed, and, therefore co-regulated (Bettencourt *et al*, 2014; Konopka *et al*., 2012) and might, thus, be functionally related (Eisen *et al*, 1998; Forabosco *et al*, 2013). It is thought that genes belonging to the same module are functionally related. This has been successfully applied to study a variety of neurodegenerative diseases (Bettencourt *et al*, 2016; Ferrari *et al*, 2016; Forabosco *et al*., 2013; Mencacci *et al*, 2015). We analyzed publicly available gene expression data from 13 regions from 47 healthy individuals constituting the GTEX v6 dataset (Consortium, 2015).

First, we co-clustered *TAX1BP1*, *ADAMTS19*, and lists of genes encoding proteins involved in lipid homeostasis, intrinsically disordered proteins, and mitochondrial proteins (table S3). We also used lists of genes in which mutations or risk variants cause PD, amyotrophic lateral sclerosis (ALS) or Alzheimer’s disease (AD) (table S3). We identified several modules in which genes from more than one of those lists cluster together, suggesting that they are functionally related (table S4).

We identified the regions of the brain in which *TAX1BP1*, *ADAMTS19* and *SNCA* are expressed (table S3), as well as the modules in which they belong. We then identified the most important genes (“hub” genes) within each module based on the module membership (MM) values (table S5). *TAX1BP1* had a MM value above the 90^th^ percentile in dark magenta module in the amygdala, blue module in the caudate, green module in the cerebellum, plum1 module in the cortex, medium purple 3 module in the hippocampus and ivory module in the nucleus accumbens. Functional annotation clustering showed that the *TAX1BP1*-containing modules in which it is the hub gene are enriched for GO biological functions related to RNA processing, protein modification by small molecule conjugation, cellular localization, protein transport, cellular catabolic process and peptide biosynthetic process.

To assess the importance of various classes of genes within the modules in which *TAX1BP1*, *ADAMTS19* and *SNCA* belong, we undertook chi2 tests to determine whether they are enriched in genes encoding proteins involved in lipid homeostasis, intrinsically disordered proteins and mitochondrial proteins. We also used lists of genes in which mutations or risk variants cause PD, ALS or AD. After correction for multiple testing, we found that mitochondrial genes preferentially cluster in the blue module of the caudate, the medium purple 3 module of the hippocampus, the ivory module of the nucleus accumbens (all three of which contain *TAX1BP1*) (fig. 7A,B,C), and the blue module of the amygdala (containing *SNCA*). The blue module of the amygdala was also enriched in genes encoding proteins involved in lipid homeostasis, and the turquoise module of the anterior cingulate cortex in gene encoding intrinsically disordered proteins (both of which containing *SNCA*). These results indicate that the function of TAX1BP1 could be related to mitochondrial physiology, which is in keeping with its known role in mitophagy (Lazarou *et al*., 2015). They also indicate that the function of αSyn is related to lipid homeostasis, mitochondria and intrinsically disordered proteins, which is also in agreement with established knowledge on αSyn being a membrane/lipid-associated protein (Fanning *et al*., 2020), being involved in dysregulation of mitochondrial bioenergetics (Choi *et al*., 2022; Ludtmann *et al*., 2016; Ludtmann *et al*., 2018) and being an intrinsically disordered protein.

**Figure 7:**
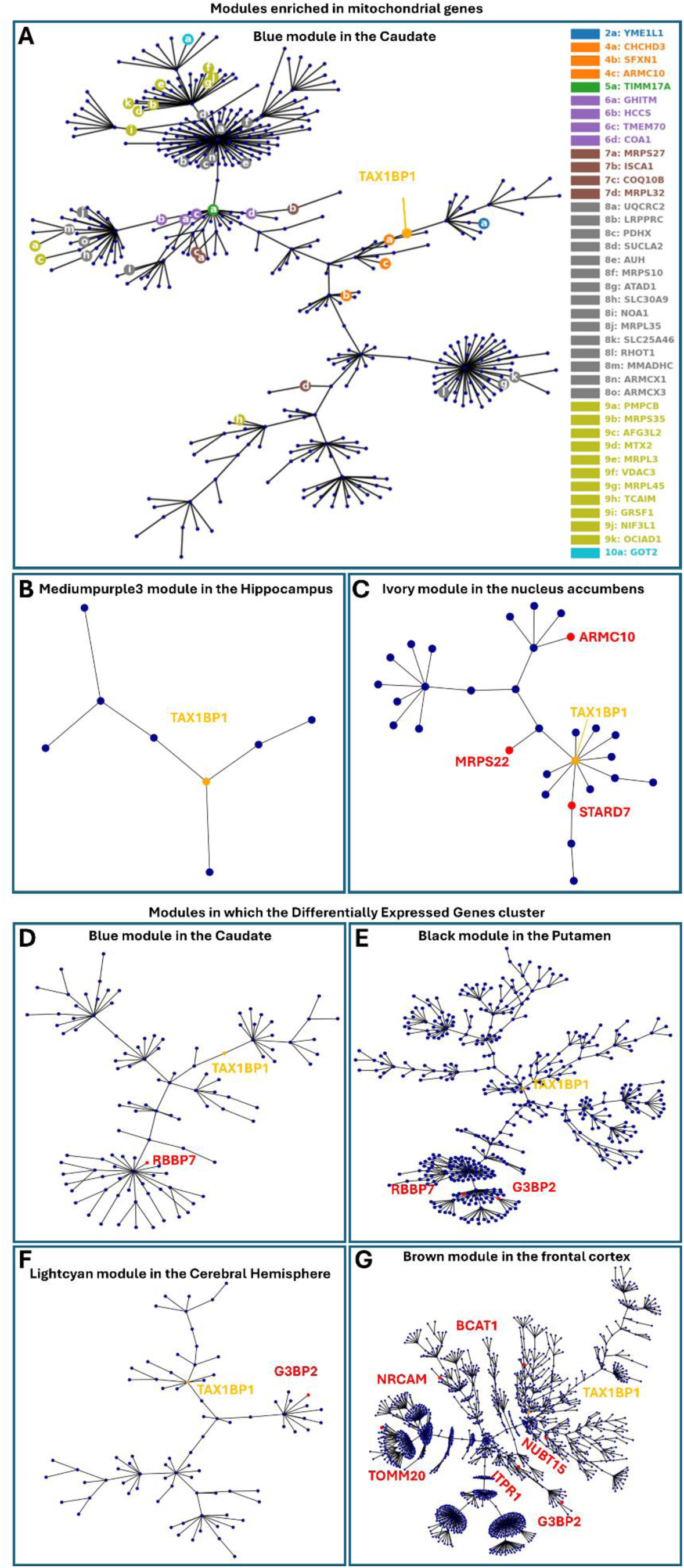
*TAX1BP1* genetic networks. A. Gene expression modules containing *TAX1BP1* that are enriched in mitochondrial genes. Blue module in the caudate. B. Medium purple 3 module in the hippocampus. C. Ivory module in the nucleus accumbens. D. Gene expression networks in which differentially expressed genes, as determined through RNA sequencing after *TAX1BP1* knock down, cluster more significantly than expected by random chance. Blue module in the caudate. E. Black module in the putamen. F. Light cyan module in the cerebral hemisphere. G. Brown module in the frontal cortex.

We then focused on the modules in which *TAX1BP1* is a “hub” gene and identified its neighboring “hub” genes, which are the genes that are highly connected within the particular module and to which it is most closely related to, indicating a close functional relevance. Those included several genes mutated or carrying risk variants in neurological diseases: *TFG* (hereditary spastic paraplegia 57) (Slosarek *et al*, 2018) in the blue module in the caudate, *CAPRIN1* (neurodevelopmental disorder) (Pavinato *et al*, 2023), *EPRS* (leukodystrophy) (Mendes *et al*, 2018) and *TFG* (hereditary spastic paraplegia 57) in the dark magenta module of the amygdala, *UBA5* (spinocerebellar ataxia) (Duan *et al*, 2016) in the ivory module in the nucleus accumbens and *CAPRIN1* (neurodevelopmental disorder) in the medium purple 3 module in the hippocampus. Of note, some of those genes neighbor *TAX1BP1* in two modules: *CAPRIN1* and *TFG*. Additionally, *TAX1BP1* neighbors to *ATXN1* and *VAMP4* within the brown module in the frontal cortex; *ATXN1* carries repeat expansions in spinocerebellar ataxia (Banfi *et al*, 1994; Orr *et al*, 1993) and *VAMP4* is a risk locus for and PD (Brown *et al*, 2021; Nalls *et al*, 2019). Finally, *TAX1BP1* neighbors *UBQLN1* in the light cyan module of cerebral hemisphere, rare variants in which cause Brown–Vialetto–Van Laere syndrome, an atypical form of motor neuron disease (Gonzalez-Perez *et al*, 2012). The fact that *TAX1BP1* is a neighbor of multiple genes involved in neurological/neurodegenerative diseases strongly supports that *TAX1BP1* has a role in the pathogenesis of neurodegeneration.

We undertook RNA sequencing after knocking down *TAX1BP1* and *ADAMTS19* in the 3K M17D cells, as described in the first section of the results (Table S1). We identified the top 30 differentially expressed genes (DEGs) for each of those conditions, as ranked by adjusted p-values, in comparison to the scrambled control. We then undertook burden analyses on whole genome sequencing data from two separate cohorts of patients and controls: 2254 sporadic cases and 2834 controls in the AMP-PD cohort, and 11841 sporadic cases and 7596 controls in the GP2 cohort, all of European ancestry. Burden analysis showed that each of those two lists of DEGs was significantly enriched in rare variants in patients with PD versus healthy controls. This was independently replicated in both cohorts (nominal p values: *ADAMTS19* DEGs AMP-PD 0.0348, GP2 0.0079; *TAX1BP1* DEGs AMP-PD 0.0044, GP2 0.0119). Those results were sub significant after application of Bonferroni correction (significance level alpha=0.0014), considering a total of 36 burden tests. Other gene lists analyzed showed occasional nominally significant p-values only in the GP2 cohort, but those were not replicated in the AMP-PD cohort (table S6). To determine whether the DEGs belong to common gene expression networks in the brain, we used WGCNA. We identified several modules in which the DEGs cluster more frequently than expected by random chance (fig. 7D,E,F,G) (table S7). Those were specific to brain regions that develop high degrees of αSyn Lewy Body pathology in PD (caudate, putamen, cortex). They were also enriched in GO functions including protein modification by small protein conjugation, protein localization, organelle organization, protein localization and endomembrane system organization. We also found that all apart from three modules were enriched for expression signatures specific to neurons. Of note, *TAX1BP1* was a “hub” gene in one of those modules: the blue module in the caudate. Based on those findings, we propose a novel type of genetic architecture for sporadic PD: an increased burden of rare risk variants across gene expression networks dysregulate the entire network and impair a particular cellular pathway, thereby leading to the development of PD.

We assessed whether *TAX1BP1* and *ADAMTS19* are differentially expressed in RNA sequencing datasets from blood from patients with PD versus healthy controls. We found that the expression of *TAX1BP1* is higher in patients with PD versus controls (p-value 0.0054) but there was no difference for *ADAMTS19* (p-value 0.21). These findings support that *TAX1BP1* is indeed dysregulated in PD, albeit the interpretation of the directionality of the change is not straightforward, given that we have found reduced expression to be associated with αSyn dysregulation in our experiments.

## DISCUSSION

It is well established that αSyn has a critical role in the pathogenesis of PD, which is supported by the presence of Mendelian mutations (Polymeropoulos *et al*, 1997) and risk variants (Nalls *et al*., 2019) in the *SNCA* gene and by the formation of αSyn-rich Lewy Bodies in neurons in the brains of patients with PD (Braak *et al*, 2003). Despite decades of efforts, the mechanism resulting in αSyn dysregulation, inclusion formation and neurodegeneration is unclear. Through a high throughput screen, we showed that knock-down of *TAX1BP1* or *ADAMTS19* increases αSyn propagation (Kara *et al*., 2021). In this work, we focused on developing a deeper mechanistic understanding of those genes.

To study their mechanism of action, we used an M17D cell line expressing 3K mutant αSyn as a model system because it recapitulates key ultrastructural features of Lewy Bodies, including entrapment of vesicles and lipids. The original report postulated that altering the KTKEGV motifs disrupts the αSyn tetramer (Bartels *et al*, 2011; Dettmer *et al*., 2015; Dettmer *et al*., 2017), thereby increasing the ratio of disordered monomers:tetramers and initiating the cascade for disease causation and pathology formation, though this is considered controversial by some (reviewed in (Lashuel *et al*, 2013)). Our findings indicate that phase separation and altered interaction with lipids are involved. First, we showed that 3K αSyn inclusions are liquid condensates that form through intermixed steps of phase separation and aggregation. This is consistent with recent findings on the biogenesis process for tau inclusions (Soeda *et al*, 2024). Second, αSyn condensates associate with lipid droplets within conglomerates. Lipid droplets are composed of a core of neutral lipids including triacylglycerol and sterol esters that is surrounded by a monolayer of phospholipids (Tauchi-Sato *et al*, 2002), whose surface is decorated by a plethora of proteins such as perilipins (Olzmann & Carvalho, 2019). Their close association is likely mediated through enhanced surface interactions between the phospholipids and the 3K αSyn (Rovere *et al*., 2019). Our molecular dynamic simulations confirmed that 3K αSyn has a higher propensity than wild type αSyn to interact with phospholipid bilayers, and CLEM experiments showed that amorphously aggregated αSyn coats the surface of vesicles and membranes entrapped within inclusions formed by 3K αSyn. Third, it is likely that binding of αSyn to the surface of lipid droplets, lipids, membranes and vesicles initiates its phase separation and lowers its phase separation threshold, which requires unnaturally high protein concentrations in *in vitro* systems (Hardenberg *et al*., 2021; Ray *et al*., 2020). In fact, it was recently shown that binding between αSyn, lipid membranes and VAMP2, a SNARE protein involved in synaptic vesicle physiology, initiates phase separation of αSyn (Agarwal *et al*., 2024) and that intrinsically disordered proteins undergo phase separation on the surface of lipid droplets (Kamatar *et al*, 2024). The fact that αSyn inclusions are rich in solid membranous structures is not mutually exclusive with them being liquid condensates: the membranous components could act as a “seed” initiating the phase separation of disordered monomers into inclusions.

The presence of three artificial mutations raises the question of the physiological relevance of the 3K αSyn model system. Mutant 3K αSyn results inclusions with similar ultrastructural features as Lewy Bodies formed in the human brain (αSyn inclusions entrapping vesicles, membranes and lipids); therefore, it is likely that the same processes are dysregulated in PD but in different ways. It is possible that lipid classes in neurons in the brains of patients with PD shift towards species with a higher proclivity to interact with αSyn and alter its propensity to phase separate. Indeed, alterations of the lipid microenvironment within cells impact the multimerization status of αSyn (Tripathi *et al*, 2022) and lipid class alterations have been observed by multiple studies that showed a decrease in phospholipid species (phosphatidylcholine, phosphatidylethanolamine, phosphatidylinositol) in various brain regions affected by αSyn pathology in humans (Cheng *et al*, 2011; Hattingen *et al*, 2009; Seyfried *et al*, 2018; Stoessel *et al*, 2018; Wood *et al*, 2018), as well as in rat and yeast models (Fanning *et al*, 2019).

*TAX1BP1* encodes an autophagy receptor (Padman *et al*, 2019) and is involved in PINK1/Parkin-mediated autophagy (Lazarou *et al*., 2015). Loss of function of TAX1BP1 is associated with inability to clear insoluble protein aggregates in the brain, indicating that TAX1BP1 is involved in aggrephagy (Sarraf *et al*, 2020). We found that *TAX1BP1* knock-down increases αSyn propagation, increases the number and decreases the size of αSyn inclusions and results in the formation of enlarged lipid droplets that are surrounded by a thin rim of αSyn. Inhibition of autophagic clearance of αSyn is likely not involved because the amount of αSyn per cell does not change after *TAX1BP1* knock-down. An alternative mechanism could be that *TAX1BP1* knock-down impairs lipophagy, which would be compatible with the known role of TAX1BP1 in autophagy. Given that the total amount of neutral lipids per cell is not impacted, it is possible that select lipid species are not being degraded efficiently, thereby leading to a shift of lipid classes and a predominance of species showing stronger interactions with αSyn. This could affect both how lipid droplets integrate within αSyn inclusions and the propensity of αSyn to propagate. Knock-down of *TAX1BP1* also results in mitochondrial hyperpolarization. It is likely that the impairment of mitophagy by *TAX1BP1* knock-down results in the accumulation of dysfunctional mitochondria with altered bioenergetics. Alternatively, it is possible that TAX1BP1 has a primary role in mitochondrial physiology, which is separate from its role in autophagy/mitophagy, and this is supported by our WGCNA data: *TAX1BP1* is a hub gene within gene expression modules that are significantly enriched in mitochondrial genes.

ADAMTS19 (A disintegrin and metalloproteinase with thrombospondin motifs 19) belongs to the family of zinc metalloendopeptidases comprising 19 members. Most of them have a known substrate but ADAMTS19 is an orphan enzyme (Kelwick *et al*, 2015). It was recently shown that KLK6, a serine protease that is highly expressed in the brain, activates ADAMTS19 through proteolysis. KLK6 cleaves αSyn fibrils that have an increased propensity for cell-to-cell propagation. Neurons with KLK6 loss take up αSyn fibrils more readily. The authors hypothesize that ADAMTS19 integrates into that pathway regulating αSyn turnover (Pampalakis *et al*., 2017). Our data indicate that *ADAMTS19* knock-down increases the propagation of αSyn and increases the number and decreases the size of αSyn inclusions. It also makes αSyn inclusions more liquid. It is possible that knock-down of *ADAMTS19* results in altered cleavage of αSyn, which produces species with a higher propensity to propagate and phase separate. We also found that *ADAMTS19* knock-down results in increased ATP levels in cells, suggesting that there is a concomitant wider dysregulation of energy production.

Our findings provide insight into the genetic architecture of sporadic PD. The genetic etiology of sporadic PD is a conundrum as genome wide association studies (GWAS) have indicated that risk loci explain only a small fraction of the “missing heritability” (Keller *et al*, 2012). Knock-down of *TAX1BP1* or *ADAMTS19* up-/downregulates a group of genes that carry an increased burden of rare variants in patients with PD in comparison to healthy controls. Those differentially expressed genes cluster in common gene expression modules in brain regions that develop high degrees of αSyn pathology. This progressively leads to the dysregulation of key cellular functions that are mediated by those networks and results in αSyn dyshomeostasis. Our approach indicates the promise of triangulating the results from high throughput screens with human genomic and transcriptomic data to dissect the genetic basis of complex diseases.

Finally, we assessed whether chloroquine, an antimalarial and disease-modifying anti-rheumatic drug (DMARD) has an effect on αSyn inclusions in the 3K αSyn cell line. We decided to test this drug because patients with rheumatoid arthritis who receive treatment with chloroquine have a significantly lower risk for the development of PD (Paakinaho *et al*., 2022). We found that chloroquine significantly reduces the number of αSyn inclusions and increases their liquid fraction when administered at 48h post-doxycycline induction, and also rescues the changes caused by knock-down of *TAX1BP1* or *ADAMTS19*. In contrast, when it is administered simultaneously with doxycycline induction, it has the opposite effect and increases the number and decreases the liquid fraction of αSyn inclusions, thereby making them more solid. Therefore, it seems that the effect of chloroquine depends on the timing of its administration in relation to the timeline in the biogenesis cycle of the αSyn inclusions. Finally, prior treatment with Oleic Acid prevented the removal of Swiss cheese αSyn inclusions (entrapping lipid droplets) by chloroquine treatment. It is possible that the interaction between αSyn and the phospholipid surface of the lipid droplets, or the change in material properties of the αSyn inclusions caused by the embedding of lipid droplets prevents penetration of chloroquine into inclusions or masks the sites of αSyn that interact with chloroquine.

## METHODS

### Tissue culture

An M17D neuroblastoma cell line stably expressing 3K mutant αSyn tagged with Venus YFP under a doxycycline induction was used (Dettmer *et al*., 2017). Those cells were maintained in medium containing Dulbecco’s Modified Eagle Medium (DMEM) with phenol red (#11965118, Thermofisher) plus 10% Fetal Bovine Serum (FBS) (#16000044, Thermofisher), 1% glutamax (#35050061, Thermofisher) and 1% Penicillin-Streptomycin ((10,000 U/mL stock), #15140122, Thermofisher) at 37°C, 5% CO_2_, and >90% humidity and were passaged at 80% confluency with Trypsin (#25300054, Thermofisher). For all the experiments, the same medium preparation was used with the exception of DMEM which did not contain phenol (#31053028, Thermofisher) to reduce background fluorescence in live imaging experiments. The cells were tested regularly for mycoplasma contamination (#NC1967484, Invivogen).

For imaging experiments, cells were seeded at a density of 2.2×10^5^ cells/mL, 300uL/well, in 8 well coverglass bottom chamberslides (#80807, Ibidi). 24h later, 3K αSyn expression was induced by treatment with 1ug/mL doxycycline hyclate (#D9891-1G, Sigma) for 48h in media containing DMEM without phenol plus 10% FBS, 1% glutamax and 1% penstrep.

Transfections were undertaken at 48h after cell seeding/24h after doxycycline induction. Two tubes were prepared: One tube containing 18uL of OMEM without phenol (#11058021, Thermofisher) plus 1.08uL of RNAiMAX (#13778150, Thermofisher) per well, and one tube containing 18uL of OMEM without phenol plus 0.3ul of pooled siRNA (5uM) and/or 240ng of construct per well. 36uL of transfection mix was added per well. Imaging experiments were undertaken 24-48h later. Cy3-tagged siRNA (#SIC003-1NMOL, Sigma) was used as a positive control for transfection.

#### Chloroquine treatment

Cells were treated with 50uM of Chloroquine diphosphate salt (#C6628-25G, Sigma) in DMEM without phenol plus 10% FBS, 1% glutamax and 1% penstrep at 48h post-doxycycline induction for 2-5h. Cells were live imaged in media containing chloroquine. In an alternative version of this experiment, cells were induced with doxycycline simultaneously with chloroquine treatment and were imaged 16h later, after replacement of the media with media of the same composition also containing Hoechst 33342 trihydrochloride trihydrate nuclear dye (#H3570, Thermofisher) (1ug/mL).

#### Spermine tetrahydrochloride treatment

Cells were treated with a dose-response of Spermine tetrahydrochloride (#S2876-5G, Sigma) concentrations (250, 500, 1000, 2000, 4000, 8000, 16000uM) in DMEM without phenol plus 10% FBS, 1% glutamax and 1% penstrep.

#### 1,6-Hexanediol treatment

Stock 1,6-Hexanediol (#240117-50G, Sigma) was warmed in a 45°C water bath until melted and formed a viscous clear liquid. Equal volumes of 1,6-Hexanediol and sterile distilled water (#15230162, Gibco) were mixed to prepare a 50%v/v stock solution (approximately 4000mM). For the experiments, final concentrations of 0.5, 1.5, 2 and 5% in DMEM without phenol plus 10% FBS, 1% glutamax and 1% penstrep were used. Cells were treated for 30min before imaging.

### Lipid droplet experiments

Treatment with pre-conjugated Oleic Acid-Albumin (#O3008-5ML, Sigma) was used to induce lipid droplet formation in cultured cells (Papadopoulos *et al*., 2015). Oleic Acid was added at 48h after doxycycline for 16h. Depending on the experiment, either a single concentration of 600uM or a dose-response (18.75, 37.5, 75, 150, 300, 600, 1200uM) were used. The cells were then fixed with 4% paraformaldehyde (PFA) (#15714-S, Electron Microscopy Sciences) and stained with Hoechst (1:2000) and HCS LipidTOX™ Deep Red Neutral Lipid Stain (#H34477, Thermofisher) (1:500). The cells were imaged on a Zeiss 780 LSM confocal microscope with a 63x oil immersion objective. The Hoechst was excited using the 405nm laser and emission was collected from 410nm to 451nm. The LipidTOX stain was excited using the 633nm laser and emission was collected from 638nm to 755nm. The Venus YFP was excited using the 488nm laser and emission was collected from 490nm to 561nm. 8 representative images were taken for each condition.

#### Chloroquine and Oleic Acid-Albumin treatments

Cells were treated with Oleic Acid-Albumin for 16h at 48h after doxycycline induction, followed by 50uM of chloroquine treatment for 5h. Afterwards, 1:2000 Hoechst was added, and cells were imaged live as described above. 8 representative images were taken for each condition.

#### Transfections

Cells were transfected as described above. At 24h post-transfection, cells were treated with 600uM of Oleic Acid-Albumin for 16h before fixing with 4% PFA and staining with Hoechst (1:2000) and LipidTOX far red neutral lipid stain (1:500). Imaging was done as described above, and 8 representative images were taken for each condition.

#### Spermine tetrahydrochloride treatment

Cells were treated simultaneously with 600uM of Oleic Acid-Albumin and 500, 1000 or 4000uM of Spermine tetrahydrochloride for 16h at 48h post-doxycycline induction.

#### 1,6-Hexanediol treatment

Cells were treated with 600uM of Oleic Acid-Albumin at 48h after Doxycycline induction, followed by 0.5, 1.5 or 5%v/v 1,6-Hexanediol treatment for 30min. Cells were fixed, stained and imaged as described above.

#### Data analysis in Python

Solid, Swiss cheese and hollow inclusions were determined as follows: Segmented inclusions were labelled with SciKit image, and individual inclusions were assessed for the presence of circular holes. Contours in the mask were extracted using contour detection (findContours() function) in OpenCV. The circularity index of the contours was computed as per the formula below, and inclusions surrounding a hole with a circularity index greater than 0.65 were deemed to have a circular inclusion (corresponding to a single, round lipid droplet) and the ones with a circularity index below 0.65 were deemed to have a clump of lipid droplets entrapped, assuming intersection between the masks for the green and red channels in those regions. Inclusions without holes were classified as solid.

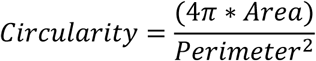

The overlap between the green and red channels was found by intersecting both Boolean matrices via binary join to identify pixels where both images were masked, showing lipid-inclusion colocalization. The surface area of the intersection of both channels was measured as the number of pixels covered by the mask. Afterwards, we divided the surface area of the intersection mask to the surface areas of the red and green masks to determine what proportion of the lipid droplets are associated with αSyn inclusions and what proportion of the inclusions are associated with lipid droplets, respectively.

### iPSc-neurons

#### Human iPS cells (hiPSC) culture

hiPSCs were incubated at 37 °C /5% CO2 incubator. Medium was exchanged every two days with 2ml mTeSR^TM^ Plus media (STEMCELL Techology, # 100-0276). Cells were passaged when colonies became ∼70% confluent (at approximately 3-5 days) with ReLeSR^TM^ (STEMCELL Techology, # 100-0483) according to manufacturer’s instructions.

#### Generation of iPSC-derived midbrain dopaminergic (mDA) neurons

Neuronal differentiation was induced as described previously (Kim *et al*, 2021b; Kriks *et al*, 2011). In detail, prior to beginning differentiation, Matrigel matrix (Corning, #354277) was thawed at 4°C overnight or on ice until dissolved. Matrigel-coated 6-well plates were prepared according to manufacturer’s instructions and incubated at 37 °C for at least 1 hour before use. hiPSCs were dissociated into single cells with Accutase (STEMCELL Technologies, # 07920) and plated at 35 × 10^4^ - 40 × 10^4^ cells per cm^2^ onto Matrigel coated plate with N2 (Neurobasal/N2 with B27 Plus Supplement; Gibco) media containing 2 mM L-glutamine (L-Glu), recombinant mouse sonic hedgehog N-terminus SHH C25II (500 ng/ml; Peprotech # 100-45), BMP inhibitor LDN 193189 (250 nM; Biogems # 1062443), ALK inhibitor SB431542 (10 μM; Biogems # 3014193), GSK3 inhibitor CHIR99021 (0.7 μM; Biogems # 2520691), and Rock inhibitor (10 μM; Y-27632, R&D systems #1254) on day 0 of differentiation, and maintained until day 3 without Rock inhibitor from day 1. On day 4, cells were exposure to 6 μM of CHIR99021 until day 10. On day 7, LDN, SB, and SHH were withdrawn. On day 10, media was changed to Neurobasal/B27/L-Glu media supplemented with 3 μM of CHIR99021 (until day 11), brain-derived neurotrophic factor (BDNF, 20 ng/ml; PeproTech # 450-02), ascorbic acid (0.2 mM; Sigma #4034), glial cell line-derived neurotrophic factor (GDNF, 20 ng/ml; PeproTech # 450-10), dibutyryl cAMP (0.2 mM; Biogems # 1698950) and transforming growth factor type β3 (TGFβ3, 1 ng/ml; R&D #243-B3). On day 11, cells were dissociated using Accutase and replated at high cell density (800 × 10^3^ cells/cm2) on dishes that were pre-coated with 15 μg/ml polyornithine / 1 μg/ml laminin / 2 μg/ml fibronectin in mDA differentiation medium (Neurobasal /B27/L-Glu, BDNF, ascorbic acid, GDNF, dibutyryl cAMP, and TGFβ3) until day 16 with the addition of 10 μM of DAPT (Biogems # 2088055) from day 12. On day 16, cells were dissociated and plated following the same procedure as day 11 and cultured until day 25 using mDA differentiation media. On day 25, cells were dissociated using Accutase and replated onto Matrigel-coated plates or coverslips at low cell density (200 × 10^3^ - 250 × 10^3^ cells/cm2) in mDA differentiation media to end point. Expression of dopamine (DA) neuron-specific and mature neuronal markers were confirmed by immunocytochemicalstaining for tyrosine hydroxylase (TH), LIM homeobox transcription factor 1a (Lmx1a) and forkhead transcription Foxa2.

#### Lipid Droplet Experiment

The iPSc-neurons were maintained by changing half the media every two days. Some of the wells were treated with 300µM Oleic Acid-Albumin for 16 hours. All the wells were stained with Hoechst (1:2000) and LipidTOX far red dye (1:500) before being imaged on a Zeiss 780 LSM confocal microscope with a 63x oil immersion objective. The Hoechst was excited using the 405nm laser and emission was collected from 410-451nm. The LipidTOX far red dye was excited using the 633nm laser and emission was collected from 638-755nm. 5-8 representative z-stack images were taken for each condition.

### Fluorescence Recovery after Photobleaching (FRAP)

FRAP experiments were undertaken as previously described (Wegmann *et al*, 2018; Zheng *et al*, 2011). All experiments were undertaken on a Zeiss LSM800 confocal with AiryScan with a 63x oil-immersion objective. FRAP experiments were optimized as follows: It was checked that the laser settings used in the timelapse portions of the FRAP experiment do not bleach the image by more than 10% in a time course acquisition of the same number of cycles as the actual FRAP experiment. The number of bleach events was also optimized: We generated a decay curve for the fluorescence intensity of the green channel by introducing one bleach iteration after each image in the timelapse and determining the number of iterations at which the fluorescence intensity was down to 10-20% of the pre-bleach value.

αSyn inclusions were bleached at 100% transmission with the 488nm laser for 8 iterations (approximately 2 seconds). Each FRAP time-lapse collected 5 cycles (frame time 5.08 seconds) pre-bleach and 35 post-bleach cycles. Three regions of interest (ROIs) were drawn in Zen 3.9 lite analysis software: ROI 1 = photobleached inclusion, ROI 2 = cytoplasm from within the same cell as ROI 1, ROI 3 = black background outside the cells. Time-lapse data of the fluorescence intensity for these three ROIs was entered into an excel spreadsheet. The fluorescence intensity of the background, ROI3, was subtracted from the intensity for the inclusion, ROI1, at each time point. A photobleaching rate (r) was determined through the control fluorescence intensity (ROI2) before and after photobleaching. The normalized fluorescence intensity after photobleaching for the inclusion (ROI1) was determined by subtracting ROI 3 from ROI 1 and dividing the value by r at each timepoint. This data was then fit to a monophasic exponential decay curve in Graphpad Prism and the immobile fraction (Fi) was calculated. The first timepoint of each FRAP time course experiment was also used to compute the circularity index for αSyn inclusions, as described above.

### Mitochondrial Membrane Potential Measurement

Cells were transfected and 48 later incubated for 40min with 100nM tetramethylrhodamine methyl ester (TMRM) (#T668, Thermofisher), a fluorescent lipophilic cation, prior to live imaging in the staining solution on a Zeiss LSM800 confocal with AiryScan with a 63x oil-immersion objective. This low concentration of the TMRM cation enables imaging in redistribution mode, which means that it can transfer in and out of the mitochondria depending on their membrane potential Δ_Ψ_, and its fluorescence intensity in the mitochondria is proportionate to the Δ_Ψ_. This system is suitable for both steady state and time course experiments (Esteras *et al*, 2020). Z-stacks were acquired for single timepoint experiments. For time course experiments, a single plane was imaged over a time course. In the time course experiments, 2ug/mL of oligomycin (#O4876-5MG, Sigma) were used to inhibit complex V, 5uM of rotenone (#R8875-5G, Sigma) to inhibit complex I and 1uM FCCP (#ab120081, Abcam) to completely depolarize the mitochondria. Experiments were imaged on a Zeiss LSM800 confocal with AiryScan with a 63x oil-immersion objective. Excitation was done with 561nm laser and emission was measured between 575-700nm. Laser power was adjusted to below 1% to reduce phototoxicity. For imaging analysis, the TMRM fluorescence was measured only in pixels contained within mitochondria, and therefore the mean fluorescence intensity (MFI) is independent of the mitochondrial mass (McKenzie *et al*, 2007).

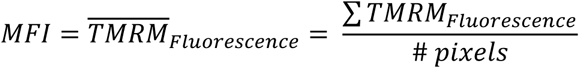

### Mitochondrial Mass Measurement

Cells were transfected and 48h later incubated for 40min with 100nM TMRM, 750nM Calcein-AM (#C3099, Thermofisher) and 1ug/mL Hoechst, followed by z-stack acquisition on a Zeiss LSM800 confocal with AiryScan with a 63x oil-immersion objective. Hoechst was excited with the 405nm laser and emission was measured between 400-464nm. Calcein was excited with 488nm laser and emission measured between 490-550nm. TMRM was excited with 561nm laser and emission measured between 575-700nm. The surface area (SA) of the mitochondria was divided to the SA of the cytoplasm, after subtracting the surface area of the nucleus for each slice of the z-stack.

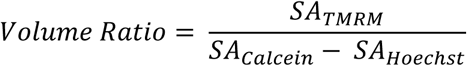

### ATP measurement

For measuring ATP levels, the construct ATeam1.03-nD/nA/pcDNA3 (Addgene #51958) was used (Kotera *et al*., 2010), according to previously described methods (Esteras *et al*, 2017). This construct contains variants of cyan fluorescent protein (CFP) and yellow fluorescent protein (YFP), which is a Forster Resonance Energy Transfer (FRET) pair. Binding of ATP to the probe brings the two fluorophores closer together and increases FRET efficiency. Therefore, FRET efficiency is proportionate to the amount of intracellular ATP levels. The cells were transfected simultaneously with the ATP construct and the siRNAs and imaged on a Zeiss 780 LSM confocal microscope with a 63x oil immersion objective 48h later. The CFP was excited using a 405nm laser and emission was collected from 460nm-499nm. Emission from the YFP was collected from 535nm-606nm. 8 representative images were collected from each well. Timelapse imaging was also performed. The CFP and FRET channels were recorded for a few minutes at baseline condition. The cells were then treated with oligomycin (2µg/mL) and immediately imaged in a 5min timelapse. Finally, cells were treated with iodoacetic acid (#I1149-5G, Sigma) (100µM) and immediately imaged with a 10min timelapse.

### Measurement of glucose uptake

Cells were transfected and 24h later treated with 250µM or 500µM of glucose fluorescent analogue 2-NBDG (2-(N-(7-Nitrobenz-2-oxa-1,3-diazol-4-yl)Amino)-2-Deoxyglucose) (#N13195, Thermofisher) in Dulbecco’s phosphate-buffered saline (DPBS) containing Calcium and Magnesium (#14040133, Gibco) for 24h, The cells were imaged on a Zeiss 780 LSM confocal microscope with a 63x oil immersion objective. The 2-NBDG was excited using the 488nm laser and emission was collected above 493nm. 8 representative images were taken from each well.

### ROS production and lipid peroxidation

#### Cytosolic ROS

Dihydroethidium (Hydroethidine) (DHE) (#D23107, Thermofisher) is a superoxide indicator that is blue fluorescent in the cytoplasm, but upon oxidization shows red fluorescence in the nucleus. 48h after transfection, the cells were treated with 10uM of DHE and timelapse imaged immediately. Cells were imaged on a Zeiss LSM800 confocal with AiryScan with a 63x oil-immersion objective. Excitation was with a 488nm lased and emission was measured between 495-700nm. 10uM of Menadione were used as a positive control (Goffart *et al*, 2021).

#### Mitochondrial ROS

Cells were treated with 100nM Mitotracker Red CM-H2XRos (#M7513, Thermofisher) (which was reconstituted under inert gas (Argon)) for 20min before doing a 10min timelapse imaging experiment. The cells were imaged on a Zeiss 780 LSM confocal microscope with a 63x oil immersion objective. Cells were excited with the 561nm laser and emission was collected from 568nm to 712nm. Two timelapses per experiment were imaged and averaged per condition.

#### Lipid peroxidation

The Bodipy 581/591 C11 dye (#D3861, Thermofisher), which is a fluorescent sensor that inserts into membranes, was used to monitor lipid peroxidation. Excitation and emission maxima are 581/591nm, which shift to 488/510nm after oxidization. 24h after transfection, the cells were treated with 2uM of Bodipy 581/591 C11 for 30min and time course imaged for 12min. Cells were imaged on a Zeiss LSM800 confocal with AiryScan with a 63x oil-immersion objective. Cells were excited using the 568nm laser and the 488nm laser and emission was collected from 574nm-650nm and 499-551nm, respectively. Two timelapses per experiment were imaged and averaged for each condition.

### Measurement of autophagy activation

For measuring autophagic flux in the cells, the construct pcDNA3-GFP-LC3-RFP-LC3ΔG (Addgene #168997) was used (Kaizuka *et al*., 2016). GFP-LC3 localizes to the autophagosome through a PE sequence and is degraded after fusion with the lysosomes, whereas RFP-LC3ΔG remains cytoplasmic. Therefore, the GFP/RFP MFI ratio can be considered a metric for autophagy activation, with lower ratios indicating higher autophagy activation. Cells were transfected with the construct and 24h later treated with 50uM of chloroquine for 2.5h prior to imaging, The cells were imaged on a Zeiss 780 LSM confocal microscope with a 63x oil immersion objective. GFP was excited using the 488nm laser and emission was collected from 493-533nm. RFP was excited using the 561nm laser and emission was collected from 566-697nm. 15 representative images were taken for each condition.

### Measurement of cell death

One hour before imaging, the media was replaced with DMEM without phenol plus 10% FBS, 1% glutamax and 1% penstrep containing the following fluorescent dyes: 20 uM propidium iodide (#P1304MP, Thermofisher) and 5ug/mL Hoechst 33342 trihydrochloride trihydrate nuclear dye (Esteras *et al*., 2017). As a positive control, cells were treated with a 5% v/v solution of 1,6-Hexanediol one hour before imaging. The cells were imaged on a Zeiss 780 LSM confocal microscope with a 63x oil-immersion objective.

### Flow cytometry

Cells were prepared for the experiment as previously described (Kara *et al*, 2017; Kara *et al*, 2018). Briefly, cells were seeded in 6 well plates at a density of 2.2×10^5^ cells/mL, induced with doxycycline 24h later, transfected 24h later, fed with DMEM w/o phenol + 10% FBS + 1% glx + 1% penstrep 24h later, and collected 24h later. For the chloroquine experiment, they were treated with 50uM of chloroquine for 5h before collection. For collection, the cells were washed with DPBS containing calcium and magnesium before adding trypsin for 1min at 37°C. The trypsin was then inactivated by adding DMEM w/o phenol + 10% FBS + 1% glx + 1% penstrep. The cells were then collected in 1.5mL Eppendorf tubes and spun down at 300g for 5min. The supernatant was removed, and the cells were resuspended in 500µL of 4% PFA for 10 minutes at room temperature before being spun down again at 300g for 5min and resuspended in 400 µL DPBS without Ca^2+^ or Mg^2+^ (#14190144, Thermofisher). The samples were stored at 4°C in the dark until flow cytometry analysis.

Flow cytometry was undertaken on the Beckman Coulter Gallios Flow Cytometer, which is equipped with 3 laser lines: 488nm, 638nm and 405nm and 10 emission filter cubes. Data was analyzed through the Kaluza 2.2 software, after gates were set based on the unstained and single-color controls.

### RNA sequencing

3K cells were seeded in 6 well plates, induced with doxycycline 24h later, transfected with siRNAs against *TAX1BP1*, *ADAMTS19* and scrambled 24h later (2 wells per siRNA). Media was replaced with DMEM without phenol + 10% FBS + 1% glutamax 24h later. 24h later (48h after transfection), the cells were trypsinized and pelleted. The two wells for the same siRNA were pooled together. Pellets were submitted to Azenta for further processing and data analysis as previously described (Chang *et al*, 2024). RNA was extracted with the PureLink RNA mini kit (#12183025, Thermofisher). The quality of the extracted RNA was assessed through Qubit RNA Assay (Invitrogen) and TapeStation analysis (Agilent). Libraries were prepared with the NEBNext Ultra RNA library prep kit (#E7770L, NEB). The trimmed reads were mapped to the reference genome using STAR aligner v.2.5.2b. Unique gene hit counts were generated through feature counts from the Subread package v.1.5.2. Differential gene expression analysis was undertaken through DESeq2. P-values and Log fold changes were generated through the Wald test.

### Molecular cloning

DNA sequences for KKTK αSyn and 3KVE αSyn fused with Venus YFP were produced through oligonucleotide synthesis and cloning into was done commercially by Twist Biosciences into the pTwist vector under the CMV promoter. Kozak sequence was inserted before the sequence of the encoded genes.

### Propagation experiment

Experiments were performed as previously described (Kara *et al*., 2021). Cells were seeded at a density of 8.7×10^5^ cells/mL in coverglass bottom chamberslides (day 1). They were transfected 3 days later (day 4) with the GFP-2A-αSyn-RFP and the GFP-2A-RFP constructs. The following day (day 5) they were treated with 50uM of chloroquine and imaged the day after (day 6). Some of the wells were treated with chloroquine on days 6 and 7. The cells were also imaged on days 7 and 8. The cells were imaged on a Zeiss 780 LSM confocal microscope with a 20x air objective. The propagation ratio was quantified in Python as per the formula below. Masks for the red and green channels were generated. The regions that were only red but not green were determined. The surface area of the red-only and the green regions was calculated, and they were divided to calculate the propagation ratio.

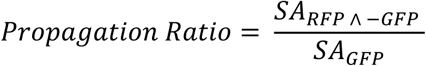

### Recombinant αSyn production

#### 3K Mutagenesis

The 3K mutant construct was created in a stepwise manner, using Q5 Mutagenesis Kit (#E0554S, New England Biolabs) according to manufacturer’s instructions. The primers used for generating the construct were designed using the NEBaseChanger tool, with sequences listed below:

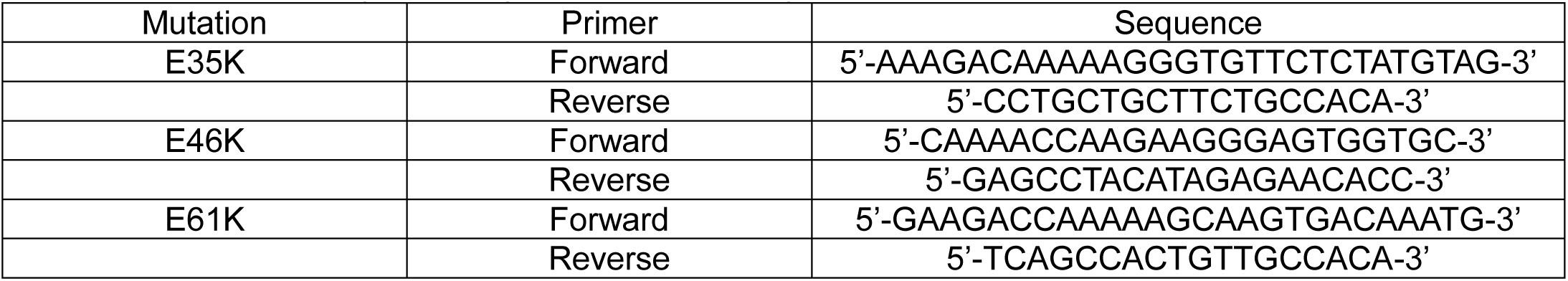

pT7-7 αSyn WT (Addgene #36046) (Paleologou *et al*, 2008) was used to make the first E35K mutation. Following the mutagenesis PCR, samples were transformed into DH5α bacteria (#C2987H, New England Biolabs) and plated on agar plates with ampicillin. Individual colonies were selected, grown, and constructs harvested for Sanger sequencing. Following confirmation of construct sequencing, the next mutation was introduced by using the pT7-7 αSyn E35K mutant plamid with the E46K primers. After sequencing, the pT7-7 αSyn E35K_E46K construct was used with the E61K primers to produce the final 3K mutant construct, followed by sequencing.

#### Mutant Protein Expression & Purification

The 3K αSyn mutant construct was transformed into BL21(DE3) bacteria (#C2527H, New England Biolabs) and plated on agar plates containing ampicillin. Starter cultures were grown by selecting colonies and growing them in LB media overnight (37°C, 180RPM). ∼5-10mL of starter culture was used to inoculate 1L LB cultures the following day, which were then grown at 37°C to an O.D. of 0.6-0.8, at which point expression was induced with 1M IPTG and grown overnight (18°C, 160RPM). Cultures were pelleted the next day and pellets were stored at −80°C until ready for use.

The 3K mutant was purified by resuspending bacterial pellets in PBS, followed by sonication (Sonicator Dismembrator, Fisher Scientific) on ice at 30% amplitude 8 cycle of 15 seconds on, 30 seconds off. Samples were subsequently boiled for 20-30 minutes, followed by centrifugation to pellet debris (20,000 RPM for 30 minutes). Nucleic acids were removed by combining the supernatant with streptomycin sulfate (final concentration of 10mg/mL) before mixing at 4°C for 30 minutes, followed by centrifugation as before. Protein was precipitated by combining the supernatant with ammonium acetate (final concentration of 0.361g/mL) followed by mixing at 4°C for 1 hour before pelleting the protein by centrifugation. The final protein pellets were suspended in Buffer A (32mM Tris, pH 7.8), filtered through a 0.2µm filter, and injected onto a HiTrap Q HP anion exchange column (Cytiva). Purification was performed on an AKTA Pure FPLC (Cytiva), with purified protein eluting at 50% Buffer B (32mM Tris, 500mM NaCl, pH 7.8).

### In vitro phase separation assays

Phase separated WT and 3K αSyn were prepared by mixing 200uM αSyn (purified using the methods provided in the protein purification section above) with different concentrations of PEG-8000 (#43443-22, Alfa Aesar, US) in LLPS buffer (25mM HEPES, 150mM NaCl, pH=7.4). The condensates aqueous sample (10uL) was added to the middle of two cover glasses that were greased together to avoid evaporation. The imaging was carried out on microscope (Nikon Ti2-A, Japan) under brightfield.

### Molecular dynamic simulations

We performed Molecular Dynamics (MD) simulations of different mutants of αSyn protein interacting with lipid membranes using the coarse-grained (CG) MARTINI 3.0.0 forcefield (Souza *et al*, 2021) with the GROMACS 2023.3 package (M.J. Abraham, 2015). Wild type and 3K and 3KVE mutant αSyn and the membrane were prepared using CHARMM GUI’s Martini Maker protocol (Hsu *et al*, 2017; Jo *et al*, 2008; Qi *et al*, 2015). CG models of the αSyn mutants were created using the martinize script (Monticelli *et al*, 2008) applied to the atomistic structures of αSyn (Ulmer *et al*, 2005) and its mutants. Flat bilayer membranes were assembled from anionic palmitoyloleoylphosphatidylglycerol (POPG) lipids the coarse-grained model of which was taken from the Martini 3 parametrization (Souza *et al*., 2021).

A single protein of each mutant was added to a POPG bilayer membrane of an approximate size 30×30 nm^2^, composed of 2900 POPG lipids, surrounded by about 95000 water beads from above and below in a cubic simulations box of approximate initial height of 17 nm, as shown in Figure 1 of the supplementary text for wild type αSyn. Salt was added to water at 100 mM concentration in form of Sodium and Chloride ions.

An initial minimization with 10,000 steps with the steepest descent algorithm was applied to each POPG bilayer with a single αSyn mutant followed by 1μs of equilibration using the velocity-rescaling thermostat (Bussi *et al*, 2007) and a Berendsen barostat (Berendsen, 1995). The production runs then followed in the isothermal-isobaric (NPT) ensemble at constant pressure, 1 bar, and fixed temperature, 303 K. Pressure coupling was applied semi-isotropically with the xy- and z-dimensions independently coupled. The equations of motions were integrated using a 20fs time step for a total simulation time of 8μs, the last 6μs of which was sampled every 6ns to calculate the average physical quantities. The production runs were simulated using the velocity-rescaling thermostat and the Parrinello-Rahman barostat (Parrinello M., 1981) with time constants of 1ps and 12ps, respectively. Simulations were analyzed with GROMACS 2023.3 built-in utilities and figures were generated using MATLAB (R2023b) and OVITO (A., 2010).

### RNA sequencing on blood samples

RNA sequencing data were taken at baseline visits from the Parkinson’s Progression Marker initiative and the Parkinson’s disease biomarkers program. We prioritized keeping features that were available for at least 80% of the training (PPMI) and validation (PDBP) cohorts available on the AMP PD platform. While RNA signatures are subject to change depending on disease stage, RNA sequencing at baseline was chosen for this analysis as it is the closest time point to diagnosis as possible. PPMI was chosen to be the training cohort given the recruitment design, recruiting unmedicated individuals within 1 year of diagnosis. Since our model is retrospective, we aimed only to analyze refined PD diagnosis, by excluding any samples with conflicting diagnostic data within a decade of post-enrollment follow-up. We excluded any cases whose medical history included an additional neurological disease diagnosis or retraction of their PD diagnosis during follow-up. We also excluded controls developing PD or another neurodegenerative disease(s) after enrollment. We included cases with clinical, genetics and transcriptomics data, including 804 PD patients and 442 healthy subjects from PDBP, and 427 PD patients and 171 healthy subjects from PPMI.

RNA sequencing data from whole blood was generated by the Translational Genomics Research Institute team using standard protocols for the Illumina NovaSeq technology. Variance stabilized counts were adjusted for experimental covariates using standard limma pipelines. Gene expression counts for protein-coding genes were extracted, then differential expression *p* values were calculated between cases and controls using logistic regression adjusted for additional covariates of sex, plate, age, ten principal components, and percentage usable bases. For RNA sequencing data, all protein-coding genes’ read counts per sample were used to generate a second set of ten principal components. All potential features representing genetic variants (in the form of minor allele dosages) from sequencing were then adjusted for the DNA sequence-derived principal components using linear regression, extracting the residual variation. This way, we statistically account for latent population substructure and experimental covariates at the feature level to increase generalizability across heterogeneous datasets. In its simplest terms, all transcriptomic data were corrected for possible confounders.

After adjustment, all continuous features were then Z-transformed to have a mean of 0 and a standard deviation of 1 to keep all features on the same numeric scale when possible. Once feature adjustment and normalization were complete, internal feature selection was carried out in the PPMI training dataset using decision trees (extraTrees Classifier) to identify features contributing information content to the model while reducing the potential for overfitting prior to model generation. We performed automated ML using the GenoML package (https://genoml.com/) on multimodal data from the PPMI. After selecting the best performing algorithm, all PPMI data was used to tune the selected model. The model was validated in the PDBP dataset.

#### Study Overview for the Parkinson’s Progression Marker Initiative

The Parkinson’s Progression Markers Initiative (PPMI), sponsored by the Michael J. Fox Foundation, is a longitudinal, observational study involving participants from 33 clinical sites worldwide. Participants contribute clinical, demographic, and imaging data, along with biological samples for whole-genome sequencing, whole blood RNA sequencing, and other analyses. The study tracks participants for a period ranging from five to 13 years. For this manuscript, we focus solely on the baseline data. This information is now available as part of AMP PD’s version 1 release. For more details about the PPMI study, please visit this link https://www.ppmi-info.org/.

#### Study Overview for the Parkinson’s Disease Biomarkers Program (PDBP)

The Parkinson’s Disease Biomarkers Program (PDBP) is an initiative led by the National Institute of Neurological Disorders and Stroke (NINDS). This longitudinal, observational study allows participants to provide clinical, demographic, and imaging data, along with biological samples for whole-genome sequencing, whole blood RNA sequencing, and other analyses. The primary objective of PDBP is to accelerate the identification of new diagnostic and progression biomarkers for Parkinson’s Disease. The data collected is now available as part of the AMP PD data releases. For further details about the PDBP study, please visit https://pdbp.ninds.nih.gov/.

### Burden analyses

We utilized genetic data from 2,254 unrelated cases and 2,834 controls of European ancestry in the Accelerating Medicines Partnership-Parkinson’s disease Initiative (AMP—PD, https://amp-pd.org/) and 11,841 unrelated cases and 7,596 controls of European ancestry from the Global Parkinson’s Genetics Program (GP2 release 6; https://gp2.org/)(Global Parkinson’s Genetics, 2021). Sequencing, sample, and variant-level quality control analyses for AMP-PD and GP2 data were conducted using the GenoTools pipeline and described elsewhere (https://github.com/GP2code/GenoTools)(Vitale *et al*, 2024). We included variants with a minor allele frequency of less than 0.05 and a minimum site depth of 20. We performed an optimized sequence Kernel association test (SKAT-O) to test the burden of 36 gene sets, including neighbors of *TAX1BP1, ADAMTS19, SNCA* as determined through WGCNA, and differentially expressed genes as determined through RNA sequencing after *TAX1BP1* or *ADAMTS19* knock-down. We used the RVTEST (Zhan *et al*, 2016) package for SKAT-O analysis by adjusting sex and the first 10 principal components.

### Experimental process and data analysis for Correlative Light and Electron Microscopy (CLEM) and cryo-ET experiments

#### Cell Lysis

A T75 flask of 3K M17D cells was pelleted. The pellet was suspended in 600ul of a hypotonic lysis buffer composed of 20mM Tris-Base, 10mM NaCl and 3mM MgCl_2_ that were dissolved in distilled water (final pH 7.4) and incubated on ice for 15min. The suspension was passed through a 25G needle 20 times and centrifuged at 3000rpm at 4°C for 5min. The supernatant contained the cytosolic and the pellet the nuclear fraction.

#### Vitrification of Cell Lysate

Cell lysates were first mixed with 6 nm gold fiducials (Electron Microscopy Sciences) to facilitate tilt series alignment during data processing. An aliquot of 3.5 μl of sample was applied to freshly glow-discharged copper Quantifoil R2/2 200-mesh London Finder grids (Quantifoil Micro Tools GmbH) and then plunge frozen using a Leica EM GP plunger (Leica Microsystems) operating at 20°C and 95% humidity. Front blotting was performed for 4-6 sec prior to plunging the EM grid in liquid ethane.

#### Correlative Light and Electron Microscopy

Cell lysate grids were loaded into a Leica DM6 FS microscope equipped with a 50x objective and a DFC 365 FX camera. The microscope stage was kept at −195°C with LN2 vapor. Fluorescence and corresponding bright field montage images of the grids were acquired using the Leica LASX software. Areas of the grid with GFP+ densities were marked as targets. Pre-screened cell lysate grids were then loaded into a 200kV Talos Arctica microscope (Thermo Fisher Scientific) equipped with a post-column BioQuantum energy filter (slit width of 20eV) and K2 direct electron detector (Gatan, Inc.). Areas of interest containing EGFP+ densities were identified at low-magnification (900x) by correlating with fluorescence images collected on Leica DM6 FS microscope. 2D projection images and tilt series of target areas were collected at 31,000x magnification with a pixel size of 4.30 Å/pixel. Imaging settings used were spot size 8, 100 μm condenser aperture, 100 μm objective aperture, and approximately −5 μm defocus. Tilt series were collected continuously from −60° to 60° with 3° step increments and in counting mode with a total dose of ∼80–100 e/Å^2^ per tilt series.

#### Cryo-ET Data Processing and Visualization

Tilt series alignment and reconstruction were done in IMOD (Mastronarde & Held, 2017). Membranes were automatically segmented using the TomoSegMemTV package (Martinez-Sanchez *et al*, 2014) and refined manually when necessary in UCSF Chimera (University of California, San Francisco) (Pettersen *et al*, 2004). αSyn aggregates, lipid droplets, and ribosomes were segmented manually using UCSF Chimera.

### Weighted Gene Co-expression Network Analysis (WGCNA)

WGCNA was undertaken on the GTEx V6 dataset for the 13 brain regions available, as previously described (Kara *et al*., 2021). Networks were generated through the WGCNA R package (Langfelder & Horvath, 2008), with an additional processing step with the k-means algorithm from the CoExpNets R package (Botia *et al*, 2017). Networks are available through the CoExp App https://rytenlab.com/coexp (Garcia-Ruiz *et al*, 2021) and also through https://github.com/juanbot/CoExpGTEx.

Clustering was undertaken for several gene lists, including known PD genes, lipid homeostasis genes, genes encoding intrinsically disordered protein, differentially expressed genes in *TAX1BP1* or *ADAMTS19* knockout, and both *TAX1BP1* and *ADAMTS19* individually. Each group of co-clustered genes were analyzed in 5 groups of brain networks, differentiated based on vulnerability to PD pathology (Braak *et al*., 2003). Clustering was considered significant if the Bonferroni corrected p value was less than 0.05. Neighboring genes were identified for *TAX1BP1*, *ADAMTS19* and *SNCA* and the module membership (MM) score was used to identify the modules in which those genes are “hub” genes. Chi2 tests were used to determine whether modules were enriched in genes belonging to certain categories.

The following gene lists were used in our analyses: Amyotrophic Lateral Sclerosis (ALS) genes: https://alsod.ac.uk/. Lipid homeostasis genes: https://maayanlab.cloud/Harmonizome/gene_set/Metabolism+of+lipids+and+lipoproteins/Reactome+Pathways Lipid genes were extracted from the reactome pathway dataset as catalogued in a Harmonizome search for genes involved in lipid metabolism or coding for lipoproteins. Mitochondrial Genes: https://www.broadinstitute.org/mitocarta/mitocarta30-inventory-mammalian-mitochondrial-proteins-and-pathways Human Mitocarta 3.0. To convert protein names to official gene symbols, each protein name was entered into the UniProt REST API search function, and the most similar response was found. Protein names with low similarity or no search results were matched by hand.

### Single Nucleotide Polymorphism (SNP) analysis

The number of single nucleotide polymorphisms was collected for each of the genes in the genome. Official gene symbols were first converted to Ensembl IDs using the Ensembl REST API. The start and end coordinate of the full gene, as well as the start and end of each exon were collected for each gene. The variation report, which contains the location of each SNP on the gene, was then collected and each SNP was identified as being located on an exon or non-exon region. SNPs were then tallied based on this distinction. The number of SNPs was normalized to the length of the exon region, non-exon region, or overall gene length to yield a frequency of variation in each region.

### Structural analyses

#### Electrostatic charges

We used PyMOL and the APBS Electrostatics Plugin to generate predictions of the electrostatic potential along the solvent accessible surface of proteins (Jurrus *et al*, 2018).

#### Disorder prediction

Metapredict was used for protein disorder prediction (Emenecker *et al*., 2021).

#### Protein charges

Charge was calculated by summing the charges of charged amino acids, Arginine (+1), Lysine (+1), Histidine (+0.1), Aspartic Acid (−1), and Glutamic Acid (−1).

### Image analysis in Python

The following general pipeline was used: .czi or .lsm files were imported in Python. Contrast was adjusted using the sigmoid, an appropriate intensity threshold was determined, and images were segmented and binarized and masks generated. For fluorescence intensity calculation within regions of interest, the masks were overlaid on the original images prior to any processing. For counting number of structures and computing surface areas, the segmented objects were labelled through functions in the SciKit library and then counted.

The Hoechst channel was analyzed as follows for counting of nuclei: Gaussian blur was first applied, followed by otsu segmentation, removal of holes and small objects with functions from the scikit-image library and object dilation with selem disk. In some analyses, the output was expressed normalized to the number of nuclei in each image. In other analyses, entire cells were segmented with the “cyto” module in Cellpose (Stringer *et al*, 2021). For specifics regarding various analyses undertaken, please see relevant sections in Methods.

### Statistical analysis

All statistical analyses were undertaken in GraphPad Prism version 10.3.0. Details on statistical tests used and number of replicates are given in the figure legends. Mean +/- Standard Deviation (SD) is shown on all the plots.

## ACKNOWLEDGMENTS

Core facilities: Confocal microscopy was performed at the school of arts and science-human genetics institute of NJ (SAS-HGINJ) Imaging Core facility and at the High Resolution Microscopy Core at the Department of Biomedical Engineering at Rutgers University. We are grateful to Drs. Zainab Tanvir and Rick Cohen, respectively, for managing those facilities. High content computing was undertaken on the Amarel cluster (Office of Advanced Research Computing – OARC) at Rutgers University. Flow cytometry was performed at the Flow Cytometry/Cell Sorting & Confocal Microscopy Core Facility at EOHSI, Rutgers University. Core operations are supported by grants from the National Institute of Environmental Health Sciences (Grant P30ES005022) and NIH (Grant S10OD026876). We thank Drs. Jessica Cervelli and Rita Hahn for core facility management.

Grants: This work was supported by Rutgers Startup funding (EK) and an HFSP long-term fellowship (LT001044/2017) (EK). NJ and WD are partially supported by NSF CAREER MCB-2046180. AHB acknowledges support from the Max Planck Society within the framework of Max Planck Partner Group and from the European Molecular Biology Organization, grant EMBO IG 5032. Supported by NIH grant NS116921 (MMM), Michael J. Fox foundation for Parkinson’s Research grant MJFF-001006 (MMM), and the American Parkinson Disease Association (MMM). ZS thank the NIH for funding this work via grants R35GM147027 and R21DA056322. JH was supported by the Dolby Family Fund.

Individuals: We are grateful to the following individuals: Jason Kaelber from the Rutgers Cryo-EM & Nanoimaging Facility (RCNF), Cynthia Pang and Sanjna Sharma for help in tomogram annotation. Dr. Patrick Lewis (Royal Veterinary College), Dr. Vikram Khurana (Harvard Medical School) and Dr. Tulsi Patel (Rutgers) for helpful discussions. Ms. Lauren Kelly for administrative assistance. Prof. Alan Goldman and Mr. Souvik Mandal for granting access to their lab for aliquoting compounds under inert gas.

NIH: This work was supported in part by the Intramural Research Program of the National Institute on Aging (NIA), and the Center for Alzheimer’s and Related Dementias, within the Intramural Research Program of the NIA and the National Institute of Neurological Disorders and Stroke. Data used in the preparation of this article were obtained from the Global Parkinson’s Genetics Program (GP2). GP2 is funded by the Aligning Science Across Parkinson’s initiative and implemented by The Michael J. Fox Foundation for Parkinson’s Research (https://gp2.org). This work used the computational resources of the National Institutes of Health high-performance computing Biowulf cluster (https://hpc.nih.gov). Data used in the preparation of this article were obtained from the Accelerating Medicines Partnership® (AMP®) Parkinson’s Disease (AMP® PD) Knowledge Platform. The AMP® PD program is a public–private partnership managed by the Foundation for the National Institutes of Health and funded by the National Institute of Neurological Disorders and Stroke (NINDS) in partnership with the Aligning Science Across Parkinson’s (ASAP) initiative; Celgene Corporation, a subsidiary of Bristol-Myers Squibb Company; GlaxoSmithKline plc (GSK); The Michael J. Fox Foundation for Parkinson’s Research; Pfizer Inc.; AbbVie Inc.; Sanofi US Services Inc.; and Verily Life Sciences. For up-to-date information on the study visit https://www.amp-pd.org.

Constructs: ATeam1.03-nD/nA/pcDNA3 was a gift from Takeharu Nagai (Addgene construct # 51958; http://n2t.net/addgene:51958; RRID:Addgene_51958). pcDNA3-GFP-LC3-RFP-LC3ΔG was a gift from Noboru Mizushima (Addgene construct # 168997; http://n2t.net/addgene:168997; RRID:Addgene_168997). pT7-7 αSyn WT was a gift from Hilal Lashuel (Addgene construct # 36046; http://n2t.net/addgene:36046; RRID:Addgene_36046).

## STATEMENT OF CONTRIBUTION

Study conception and design: EK; Study supervision: EK, JB, AHB, ZS, WD, MMM, JH; Funding acquisition: EK, JH, JB, AHB, ZS, WD, MMM; Generation of iPSc-neurons: JL; FRAP experiments and analysis: EE, SA, KV, NP, PS, EK; Electrostatic analyses: JM, EK; Wrote code for imaging analysis: JM, RK, BS, EK, AG, NP WGCNA: BS, NP, PS, JM, EK; Wrote the paper: EK, KV, EE, JM, BS, NP, NJ, WD, JW, JE, FA, SBC, FH, JL, MMM, AMB; Edited the paper: EK, JB, ZS, UD, TB, MMM, NJ, WD, JH, KV, NP; Provision of critical reagents and advice: TB, UD; Tissue culture: NP, KV, EE, EK, PS, BS, JM; Mitochondrial mass measurement: BS, PS; Mitochondrial membrane potential: BS, PS, EE, KV, RK, EK; ATP measurement: JM, KV, EK; Flow cytometry: KV, SA, EE, NP; Lipid imaging: KV, RK, NP, EK; Propagation imaging: RK, EK, SA; Measurement of cell death: EE, KV, JM, PS, EK; LC3 imaging: JM, KV, PS; ROS measurement: EE, KV, JM, PS; Glucose uptake experiments: KV; Measurement of synuclein inclusions: EK, EE, NP, KV, NP; SNP analyses: BS, JM; SNP browser development: BS; RNA sequencing: EK, EE; Experiments with recombinant synuclein: MDF, JW, JE, BS; CLEM: NJ, WD; Molecular dynamic simulations: FH, AHB; Burden analysis: FA, SBC; RNA sequencing analyses of human blood samples: SBC.

## Conflict of interest

M.M.M. is an inventor of filed and issued patents related to α-synuclein. M.M.M. is a founder of MentiNova, Inc. E.K. is a member of the EMBO Scientific Exchange Grants Advisory Board.

## DATA AVAILABILITY

Code: https://github.com/eleannakara/Kara-Lab/tree/main/synuclein_inclusion_biogenesis_2024, RNA sequencing: GEO accession numbers will be made available to reviewers and publicly upon publication of the manuscript., SNP browser: <u>karalab.shinyapps.io/SNP_Browser/</u>, Tomograms of cell lysate showing amorphous alpha-synuclein associated with vesicles and lipid droplet, and with a mitochondria outer membrane have been deposited in the EMDataBank under accession codes that will be made available to the reviewers and publicly upon publication of the manuscript.

---

**Supplementary figure 1:**
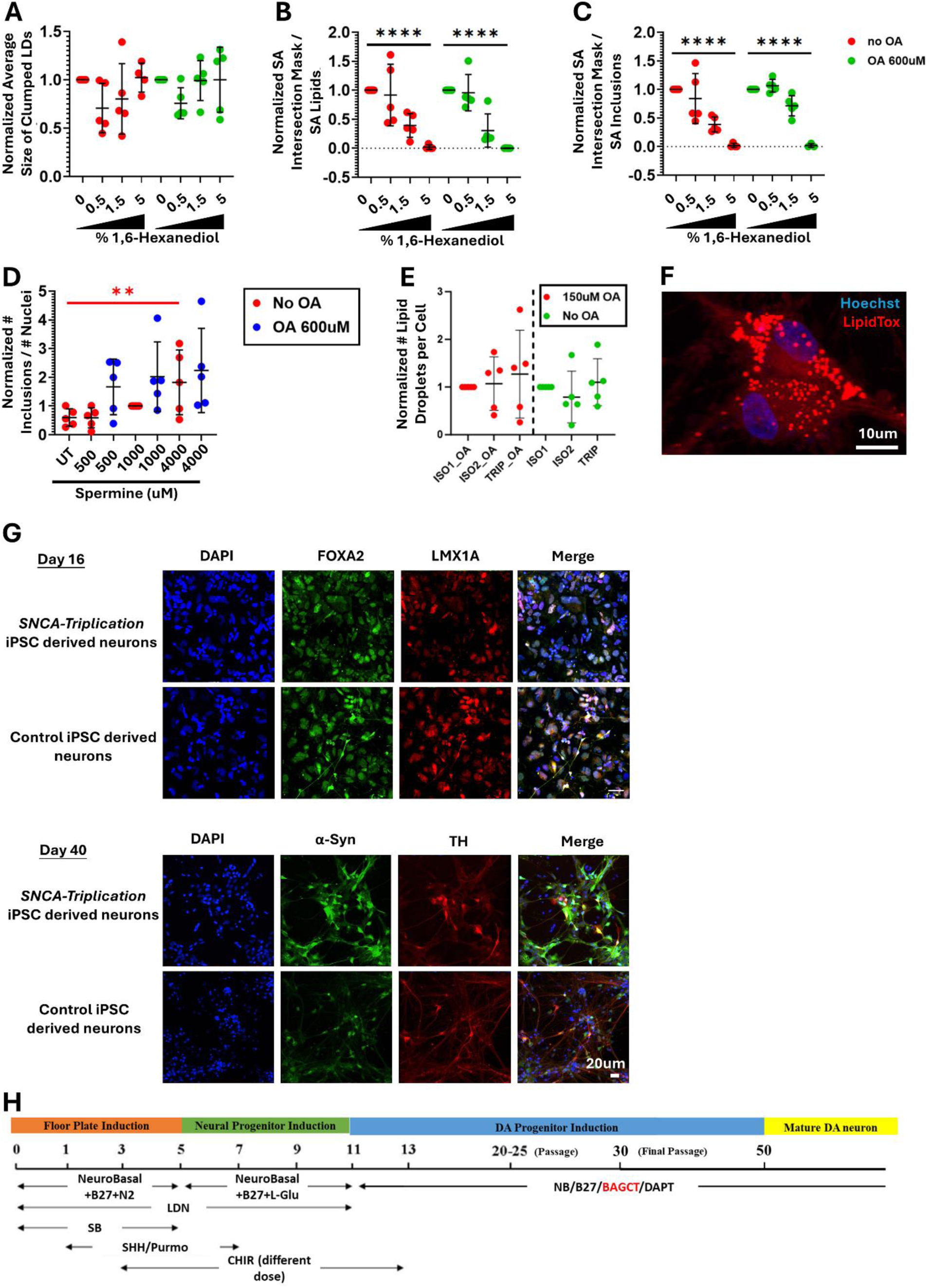
Lipid droplets in 3K αSyn cells and in dopaminergic iPSc-neurons. Related to figure 3. A. Average size of clumped lipid droplets after treatment with a dose-response of 1,6-Hexanediol concentrations (%v/v). B. Plot showing the fraction of the overlapping surface areas of the red (LipidTOX) and green (αSyn) masks relatively to the total surface area of the red mask. The results represent the fraction of the lipid droplets that are not embedded within αSyn inclusions after treatment with a dose-response of 1,6-Hexanediol concentrations (%v/v). C. Plot showing the fraction of the overlapping surface areas of the green (αSyn) and red (LipidTOX) masks relatively to the total surface area of the green mask. The results represent the fraction of the αSyn inclusions that are not associated with lipid droplets after treatment with a dose-response of 1,6-Hexanediol concentrations (%v/v). D. Number of αSyn inclusions per cell that were treated with a concentration gradient of Spermine plus 600uM of Oleic Acid for 24h. The test for trend was significant only for the condition without OA treatment. E. Dopaminergic iPSc-neurons were treated with 150uM of Oleic Acid for 16h and the number of lipid droplets was quantified. Lines included were derived from patient fibroblasts with *SNCA* triplication and two isogenic controls. One way ANOVA with Dunnett test correction for multiple testing was done separately for the OA treated and untreated cells. F. Representative confocal image of (E). G. Immunocytochemical verification of the phenotype of dopaminergic iPSc-neurons. A unique feature of the developing midbrain is co-expression of the floor plate marker FOXA2 and the roof plate marker LMX1A. H. Diagram of culture conditions for dopaminergic (DA) neurons differentiation. Neurobasal/B27 supplement (NB/B27). Purmorphamine (Purmo), brain-derived neurotrophic factor (BDNF)/ ascorbic acid/ glial cell line-derived neurotrophic factor (GDNF)/ dibutyryl cAMP/ transforming growth factor type β3 (TGFβ3) (BAGCT). For plots A-C, the data was normalized separately for the OA and non-OA treated samples to the condition without 1,6-Hexanediol. Data was analyzed through a one way ANOVA with test for linear trend, separately for the OA and the non-OA conditions. For plot D, the data was normalized to the Spermine 1000uM concentration and one way ANOVA with linear test for trend was done separately for the OA versus non-OA treated samples. 5 biological replicates were done for each experiment, unless otherwise stated. Plots containing normalized data are labelled as such on the Y axis. P-values: *≤0.05, **≤0.01, ***≤0.001, ****≤0.0001. Only statistically significant differences are shown in the plots. 1,6-Hexanediol concentrations are in %v/v. Abbreviations: LDs: lipid droplets; SA: surface area; OA: Oleic Acid.

**Supplementary figure 2:**
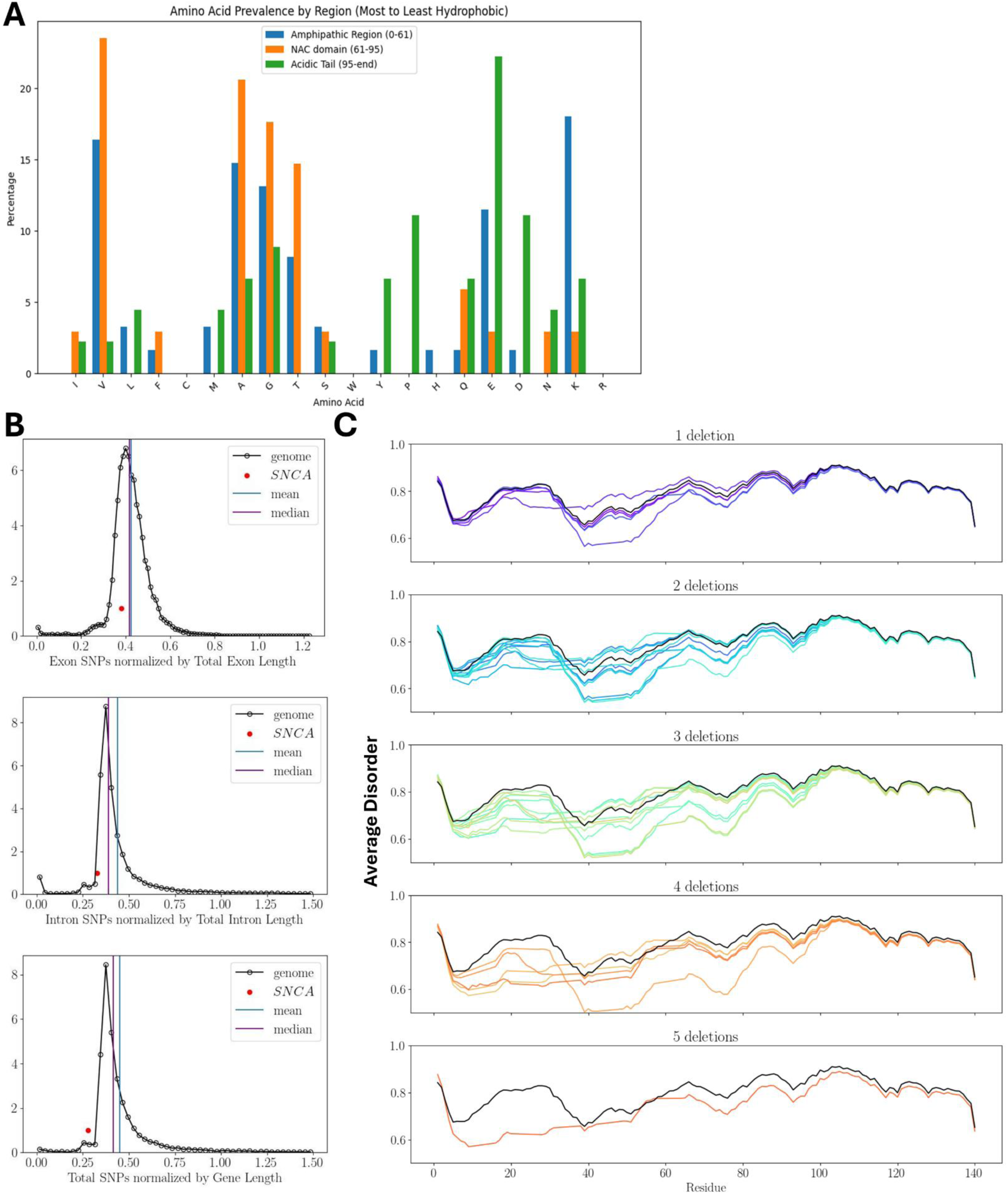
The molecular grammar of αSyn. Related to figure 4. A. Plot showing the percentage of each amino acid in each region of the αSyn protein. B. Frequency plots showing the number of single nucleotide polymorphisms (SNPs) normalized to the total length of the exons, introns or the entire gene for the *SNCA* gene in comparison to every gene in the genome. C. Average disorder of the αSyn protein following the *in silico* deletion of combinations of the first five KTKEGV repeat motifs (shown in figure 1B and 4H). The wild type αSyn protein is depicted in black. Each of the colored lines depicts one of the *in silico* generated mutants.

## TABLE LEGENDS

**Table S1:** Differentially expressed genes between scrambled siRNA and *TAX1BP1* knock-down (tab 1), and scrambled siRNA and *ADAMTS19* knock-down (tab 2). Columns E-J show normalized hit counts.

**Table S2:** Amino acid substitutions used in the *in silico* analyses shown in figure 4F,G.

**Table S3:** Weighted Gene Co-expression Network Analysis (WGCNA). Tab 1: Lists of genes used in the analyses reported in this manuscript. Tab 2: Groups of brain regions used in the WGCNA clustering. Tab 3: Genes expressed in the 13 brain regions studied, as well as in the entire brain. Tab 4: Neighbors of the *TAX1BP1* gene, along with their annotation. Tab 5: Neighbors of the *ADAMTS19* gene, along with their annotation. Tab 6: Neighbors of the *SNCA* gene, along with their annotation. Tab 7: Genes included in the modules that contain *TAX1BP1*. Tab 8: Genes included in the modules that contain *ADAMTS19*. Tab 9: Genes included in the modules that contain *SNCA*. Tab 10: Chi2 analyses to identify whether modules containing *TAX1BP1* or *SNCA* are enriched in specific categories of genes. Abbreviations: IDP: intrinsically disordered proteins.

**Table S4:** WGCNA clustering of various gene lists in combinations of brain regions. Statistically significant modules are shown (tab 1). Tab 2: Genes contained within the significant modules listed in tab 1. Column E lists the genes from the lists that cluster in the particular module. Columns M-S list how many genes from each of the lists that were clustered are present in column E.

**Table S5:** Module Membership (MM) values for the modules in which *TAX1BP1*, *ADAMTS19* or *SNCA* are present. Column G lists the 90^th^ percentile of the MM value for each module. Columns H-J list the MM values for each of the 3 genes in the respective module.

**Table S6:** Burden analyses. Neighboring genes for *ADAMTS19*, *SNCA* or *TAX1BP1*, as well as the top 30 differentially expressed genes (DEGs) (as ranked by adjusted p-values) after knock-down of *ADAMTS19* or *TAX1BP1* were analyzed.

**Table S7:** Modules in which the DEGs after knock-down of *ADAMTS19* or *TAX1BP1* cluster.

## Notes

### Competing Interest Statement

M.M.M. is an inventor of filed and issued patents related to α-synuclein. She is a founder of MentiNova, Inc.

