## Supplementary Information for "Increased burden of rare risk variants across gene expression networks predisposes to sporadic Parkinson’s disease"

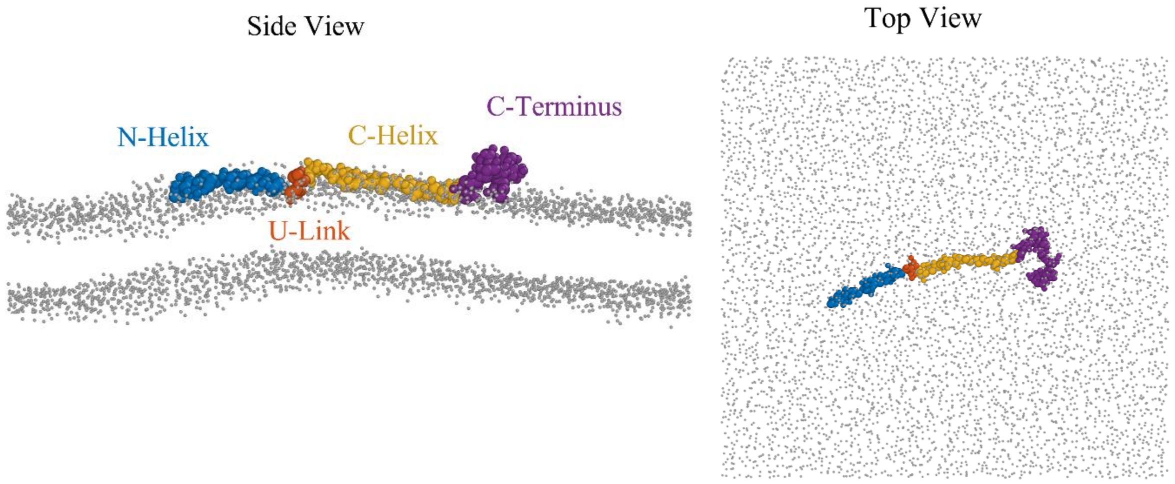


Figure 1. Side and top views of coarse-grained representation of the wild-type αSyn in the Martini model. Different regions of the protein including the N-Helix (residues 1-33), U-Link (residues 34-44), C-Helix (residues 45-92), and the C-Terminus (residues 93-140) are shown in blue, orange, yellow, and purple, respectively. The head groups of POPG lipids are shown in grey.

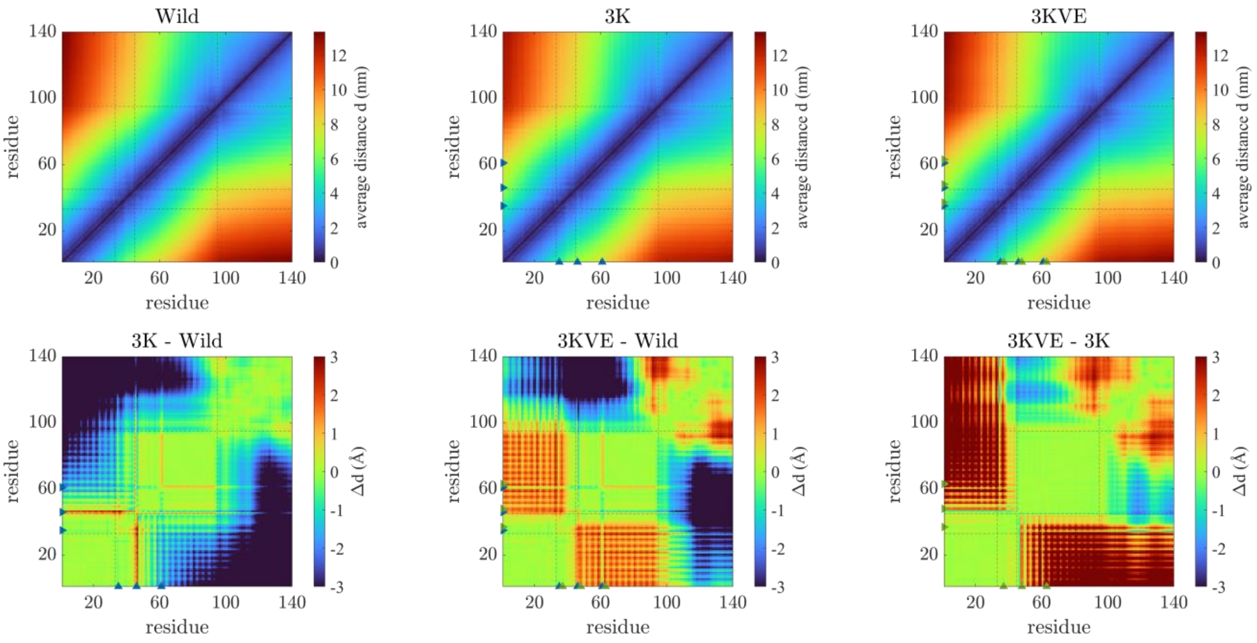


Figure 2. Conformation maps showing the distances between residue pairs in different αSyn mutants (top panels) where the average distance d is calculated between the centers of mass of the two residues. The difference maps (bottom panels), obtained by subtracting the conformation maps of the two corresponding mutants, highlight the difference between mutants. Blue triangle

markers correspond to the mutated residues 35, 46, 61, and green markers indicate the mutated residues 37, 48, 63, both with respect to the wild type αSyn.

The top panels in Figure 2 show the conformation maps of different wild-type, 3K, and 3KVE mutants of αSyn. The maps represent the average distance d between all pairs of protein residues calculated as the average of a thousand frames sampled from CG simulations of a single mutant on a POPG bilayer membrane during 6μs-long MD simulations, see Methods for more details on the simulation protocol. Each confirmation map is partitioned, by dashed lines, into four sections of N-Helix, U-link, C-Helix, and the C-terminus as shown in CG snapshots in figure 1. Also shown in figure 2 are the difference conformation maps between the mutants which better show the effect of individual mutations on an Angstrom scale as shown in the bottom panels of figure 2. Particularly, the residue 46 in the C-Helix domain of the 3K mutant exhibits an increased average distance from most of the N-Helix residues compared to the wild type mutant. The difference between the 3K and wild mutants shows that residue 46 in the 3K mutant has increased distance of about ~3 Angstroms from the N-Helix (residues 1-33) compared to the wild type. A similar difference is seen for the residue 46 between 3KVE and the wild type (see the middle bottom panel in figure 2) whereas the 3K and 3KVE mutants do not remarkably differ.

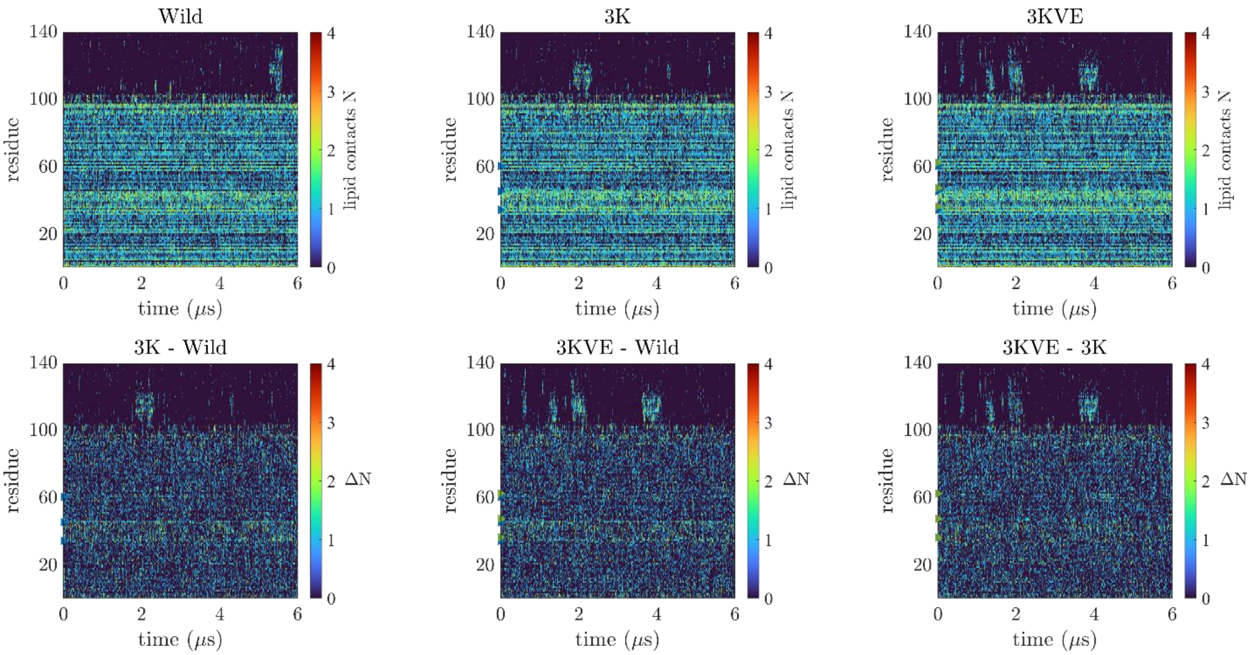


Figure 3. Contact maps showing the number of lipids N in contact with each residue as a function of time for different mutants (top panels) and the difference map where contact maps are subtracted to show differences between the mutants (bottom panels). Blue triangle markers correspond to the mutated residues 35, 46, 61, and green markers indicate the mutated residues 37, 48, 63, both with respect to the wild type αSyn.

The contact maps of αSyn mutants are shown in figure 3. These maps indicate the time evolution of number of lipids, N, in contact with each residue. A POPG lipid is considered in contact with a residue if any of the lipid head groups are located within a 0.7 nm distance from the center of mass of the residue. The lipid contacts N is proportional to the depth of the protein residue inside the lipid bilayer. Although the contact maps of the three mutants look similar, the difference maps between each two mutants exhibit difference in ΔN. The difference ΔN shows an increase at the two first mutated residues 35 and 46 between 3K and wild as well as 3KVE and wild (see the two horizontal yellow lines in the two bottom left panels in figure 3) while there is no remarkable difference between 3KVE and 3K.

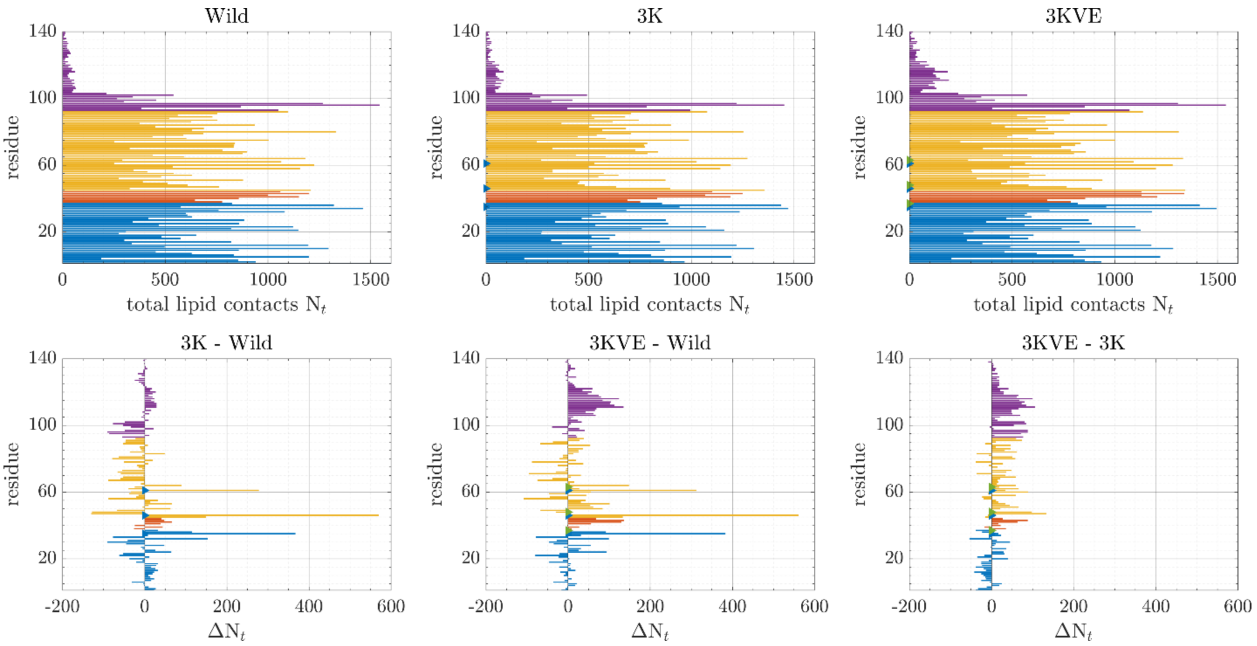


Figure 4. Accumulated contact maps showing the total number of lipids $N_t$ in contact with each residue during the entire simulation for each mutant (top panels), and the difference maps where $\Delta N_t$ is calculated by subtracting the corresponding contact maps of every two mutants (bottom panels). Blue triangle markers correspond to the mutated residues 35, 46, 61, and green markers indicate the mutated residues 37, 48, 63, both with respect to the wild type αSyn.

The accumulated contact maps in figure 4, top panels, show the total number of lipids $N_t$ whose head groups have been within a 0.7 nm distance from the centre of mass of each residue at least in one simulation frame. The difference in the accumulated contact maps (bottom panels of figure 4) exhibit three peaks at the mutated residues 35, 46, 61 between both the 3K and 3KVE mutants as compared to the wild type, implying deeper insertion of the mutated residues in 3K and 3KVE mutants inside the lipid bilayer.

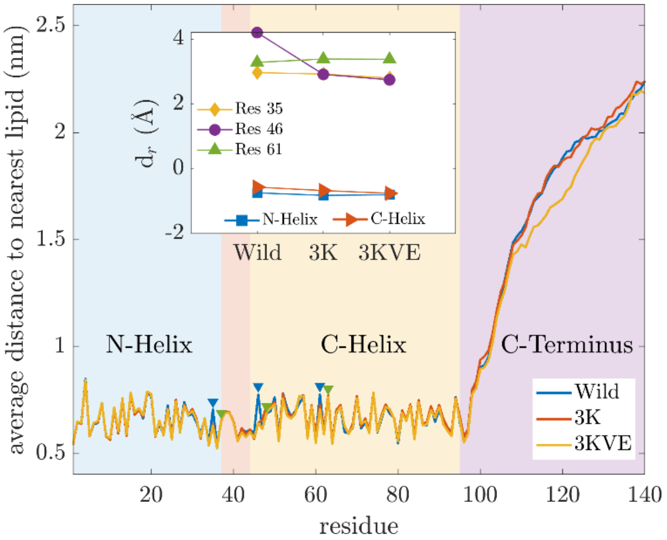


Figure 5. The average distance of each residue to the nearest lipid during the 6μs-long simulation. Blue triangle markers correspond to the mutated residues 35, 46, 61, and green markers indicate the mutated residues 37, 48, 63, both with respect to the wild type αSyn. The inset shows the relative depth $d_r$ of the mutated residues and the centres of mass of the N- and C-Helices with respect to the lipid bilayer's top surface for each mutant.

Figure 5 shows the average distance of each residue to the nearest lipid head group throughout the 6μs-long simulation. The distance to nearest lipids is reduced at the mutated residues from wild type to 3K and 3KVE mutants as seen in the difference of the blue curve from the orange and red curves in figure 5. The inset shows the relative depth $d_r$ of the mutated residues 35, 46, and 61 as well as the depth of the centre of mass of the N- and C-Helices with respect to the top surface of the lipid bilayer. The top bilayer surface is obtained by averaging over the height of the lipid head groups which are within a 1x1 $nm^2$ patch around the corresponding residue or center of mass in the top bilayer leaflet. The mutated residues 35, 46, and 61 show deeper insertion inside the lipid bilayer for 3K and 3KVE compared to the wild type, with the most significant decrease in the average distance observed at residue 46.
